# TAZ inhibits GR and coordinates hepatic glucose homeostasis in normal physiologic states

**DOI:** 10.1101/2020.05.12.091264

**Authors:** Simiao Xu, Yangyang Liu, Ruixiang Hu, Min Wang, Oliver Stöhr, Yibo Xiong, Liang Chen, Hong Kang, Lingyun Zheng, Songjie Cai, Li He, Cunchuan Wang, Kyle D. Copps, Morris F. White, Ji Miao

## Abstract

The elucidation of the mechanisms whereby the liver maintains glucose homeostasis is crucial for the understanding of physiologic and pathologic states. Here, we show a novel role of hepatic transcriptional co-activator with PDZ-binding motif (TAZ) in the inhibition of glucocorticoid receptor (GR). TAZ interacts *via* its WW domain with the ligand-binding domain of GR to limit the binding of GR to gluconeogenic gene promoters. Therefore, liver-specific TAZ knockout mice show increases in glucose production and blood glucose concentration. Conversely, the overexpression of TAZ in mouse liver reduces the binding of GR to gluconeogenic gene promoters and glucose production. Thus, our findings demonstrate distinct roles of the hippo pathway effector proteins yes-associated protein 1 (YAP) and TAZ in liver physiology: while deletion of hepatic YAP has little effect on glucose homeostasis, hepatic TAZ protein expression decreases upon fasting and coordinates gluconeogenesis in response to physiologic fasting and feeding.

## Introduction

The liver plays a critical role in organismal energy homeostasis by regulating diverse biologic processes in response to nutrient availability^1^. During fasting, the activation of hepatic gluconeogenesis is required for the supply of glucose to tissues with a high glucose demand, such as the brain, and for the maintenance of glucose homeostasis, whereas in the fed state gluconeogenesis is suppressed^2, 3^. Precise control of hepatic gluconeogenesis is crucial for normal physiology, and a failure to suppress hepatic gluconeogenesis post-prandially contributes to hyperglycemia in insulin resistance and diabetes^4^.

Glucagon and glucocorticoids (GCs; in humans, cortisol; in mice, corticosterone; synthesized in the adrenal cortex) are hormones that are secreted during fasting and promote hepatic gluconeogenesis at multiple levels, including *via* gene transcription. Glucocorticoid receptor (GR, encoded by the *NR3C1* gene) is a member of the nuclear receptor super-family^5^ that is a key transcriptional regulator of gluconeogenic gene expression in the fasting state^6^. GR not only directly responds to increases in GC concentration by activating gluconeogenic gene transcription, but also plays a permissive role in the glucagon-mediated transcriptional control of these genes. Therefore, the deletion of hepatic GR leads to fasting hypoglycemia^7^, while adrenalectomy abrogates the induction of gluconeogenesis by fasting, glucagon, cAMP, or epinephrine in rodents^8, 9^. Similarly, a single dose of a GR antagonist is sufficient to reduce hepatic glucose output in healthy humans^10^.

GR transactivation of gluconeogenic genes involves a series of molecular events^6^. Upon the binding of a ligand (a synthetic agonist or an endogenous GC)^11^, GR undergoes conformational changes, translocates to the nucleus, dimerizes, and binds to glucocorticoid response elements (GREs) in the promoters of key gluconeogenic genes, such as phosphoenolpyruvate carboxykinase 1 (*PCK1*) and glucose-6-phosphatase catalytic subunit (*G6PC*)^11, 12^, which encode the rate-limiting and final enzymes of gluconeogenesis, respectively^13, 14^. This activation mechanism differs from that of the transrepression of inflammatory genes by GR, which involves the tethering of monomeric GR to DNA-bound proinflammatory transcription factors^15^.

Yes-associated protein 1 (YAP) and transcriptional co-activator with PDZ-binding motif (TAZ) are downstream effectors of the Hippo pathway^16^. Inhibition of the Hippo pathway activates YAP and TAZ, and they co-activate TEA domain (TEAD) transcription factors within the nucleus, which induce the expression of genes involved in cellular proliferation^16^. However, YAP and TAZ also play other roles, mediated by interactions with diverse transcription factors^17^. Although YAP and TAZ share nearly 50% amino acid sequence similarity, the two proteins have distinct functions that are exerted *via* interactions with different transcription factors^17, 18^. For example, YAP activates epidermal growth factor receptor 4 (ErbB4) and tumor protein p73^19, 20^, whereas TAZ specifically interacts with peroxisome proliferator-activated receptor gamma (PPARγ) and T-box transcription factor (TBX5)^21, 22^. In addition, although TAZ co-activates TEADs, it also acts as a repressor: the binding of TAZ to PPARγ inhibits the PPARγ-induced differentiation of mesenchymal stem cells into adipocytes^21^. Thus, YAP and TAZ have distinct roles in gene promoter-specific transcriptional regulation.

We previously reported that YAP integrates gluconeogenic gene expression and cell proliferation, which contributes to its tumorigenic effects. However, YAP does not regulate normal glucose homeostasis, because hepatocyte-specific deletion of YAP in normal mice has little effect on gluconeogenic gene expression or blood glucose concentration^23^. Whereas YAP is primarily expressed in cholangiocytes in normal mouse liver^24^, we show here that TAZ is abundantly expressed in pericentral hepatocytes and that its expression is markedly reduced by fasting. In accordance with these data, we show that TAZ, but not YAP, interacts with GR to inhibit the GR-transactivation of gluconeogenic genes, thereby coordinating hepatic glucose production with physiologic fasting and feeding in normal mouse liver.

## Results

### Fasting and feeding alters hepatic TAZ protein level

To determine whether TAZ plays a role in hepatic metabolic regulation under normal physiologic conditions, we measured hepatic TAZ expression in mice that had been fed *ad libitum* or fasted for 24 h. As expected, fasting increased the hepatic mRNA expression of genes encoding the key gluconeogenic enzymes *Pck1* and *G6pc,* but not those of canonical TAZ and YAP target genes involved in cell proliferation^16^, including connective tissue growth factor (*Ctgf*) and cysteine-rich angiogenic inducer 61 (*Cyr61*) (Figure 1–Figure supplement 1A–B). Hepatic TAZ protein level was reduced by >50% after 24 h of fasting, relative to *ad libitum* feeding (Figure. 1A–B). The phosphorylation of TAZ at serine 89, which is mediated by the Hippo pathway core kinase large tumor suppressor kinase (LATS)1/2^25^, was commensurate with the reduction in total TAZ (Figure 1A–B). Both nuclear and cytoplasmic TAZ proteins shared this regulation (Figure 1C–D).

**Figure 1.**
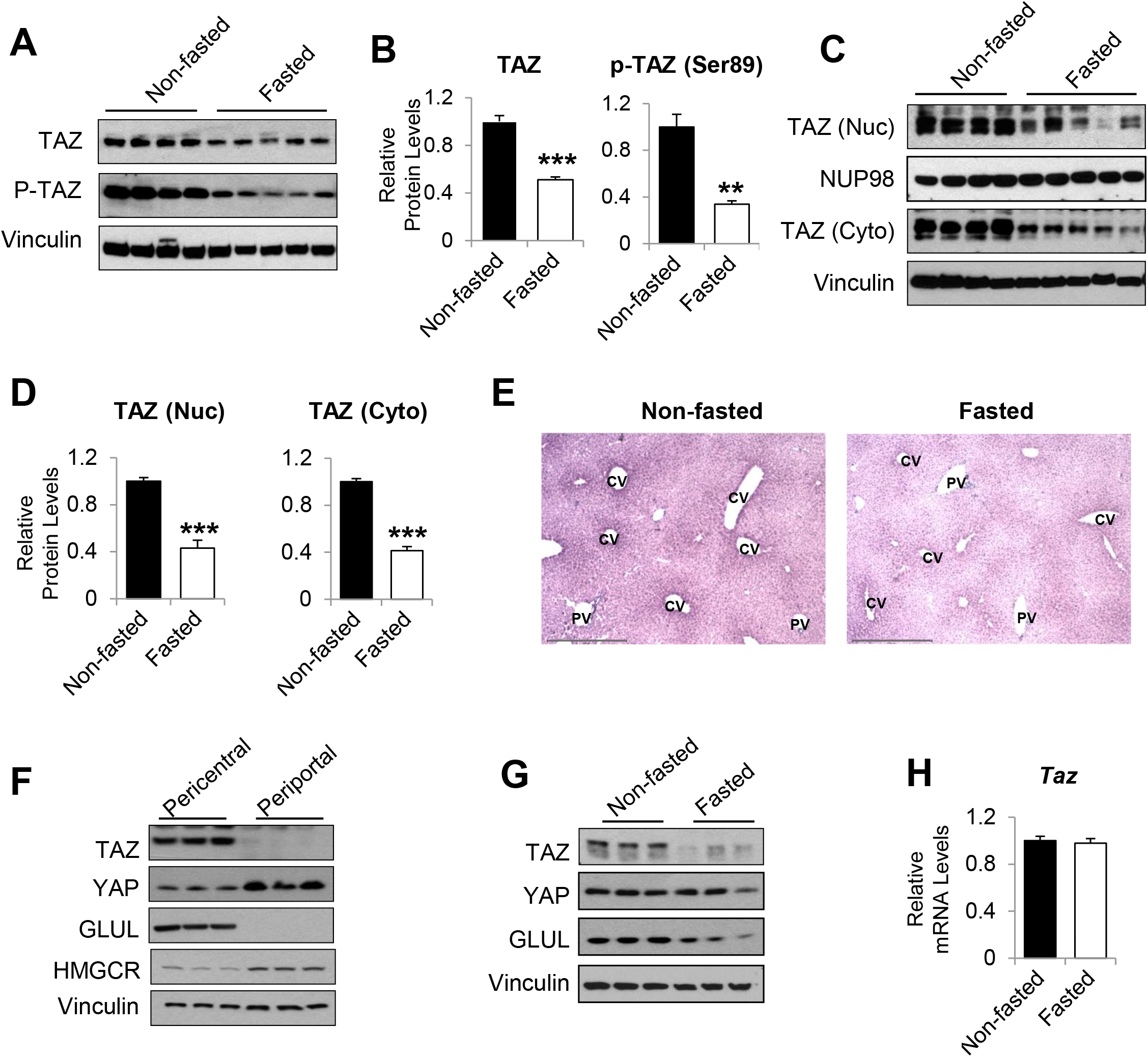
TAZ protein is regulated by fasting and feeding in hepatocytes. Eight- to twelve-week-old C57BL/6J mice were *ad libitum*-fed (Non-fasted) or fasted for 24 h (Fasted). (A–D) Hepatic proteins were measured by immunoblotting whole cell lysates (A; quantified results are shown in B) or nuclear (Nuc) or cytoplasmic (Cyto) extracts (C; quantified results are shown in D). (E) Immunohistochemical staining for TAZ in the livers of mice fed *ad libitum* or fasted for 24 h (CV, central vein; PV, portal vein; scale bar, 500 μM). (F–G) Protein levels were measured by immunoblotting lysates of pericentral and periportal mouse hepatocytes (F) or hepatocytes isolated from mice that were *ad libitum*-fed or fasted for 24 h (G). (H) mRNA expression of hepatic *Taz* was measured using real-time qRT-PCR. Data are means and SEMs; control values were set to 1; n = 6. Representative results of two to three independent experiments are shown. Data were analyzed by unpaired Student’s *t*-test; \*\**p* < 0.01 and \*\*\**p* < 0.001.

Immunohistochemistry confirmed that TAZ was abundantly expressed in both the nuclear and cytoplasmic compartments of mature hepatocytes and that its expression was reduced by fasting (Figure 1E). The antibody used was validated using liver sections from liver-specific TAZ knockout (L-TAZ KO) mice (which were generated by crossing TAZ floxed mice with Albumin-cre mice) (Figure 1–Figure supplement 2). Interestingly, TAZ showed zonal expression, with the highest protein levels being found in the pericentral (or perivenous) hepatocytes (Figure 1E). This zonal expression negatively correlated with the expression and activity of gluconeogenic genes in the liver lobule^26^. Immunoblotting of isolated pericentral and periportal hepatocytes confirmed that TAZ was primarily expressed in glutamine synthetase (GLUL)-expressing pericentral hepatocytes; whereas YAP is primarily expressed in periportal hepatocytes (Figure 1F). These results are consistent with the higher TAZ mRNA expression in pericentral than periportal mouse hepatocytes, revealed by single-cell RNA analysis^27^. In addition, TAZ protein was less abundant in hepatocytes isolated from fasting than *ad libitum*-fed mice (Figure 1G), suggesting that fasting reduces TAZ protein in hepatocytes.

In contrast to TAZ protein levels, TAZ mRNA was not affected by fasting or feeding (Figure 1H), indicating that TAZ is post-transcriptionally regulated by physiologic fasting and feeding. These data are consistent with previous findings that TAZ is subject to ubiquitin-mediated degradation^28^. Consistent with this, TAZ protein, but not its mRNA, was induced in mouse primary hepatocyte cultures by supplementation of the medium with a high glucose concentration (25 mM) and 10% fetal bovine serum (FBS), which mimics the fed condition, in a time-dependent manner (Figure 1–Figure supplement 3).

### Knockdown or knockout of hepatic TAZ induces gluconeogenic gene expression and hyperglycemia in mice

To define the role of TAZ in glucose homeostasis in mouse liver, we acutely knocked down hepatic TAZ using adenoviruses expressing AdshTAZ or control AdshCon. Compared with the control virus, administration of AdshTAZ to C57BL/6J mice reduced hepatic TAZ mRNA and protein levels (Figure 2A–B). TAZ knockdown did not alter mouse body or epididymal white adipose tissue mass, but it slightly reduced liver mass, without affecting liver histology (Figure 2–Figure supplement 1A–D). However, TAZ knockdown significantly increased the mRNA expression of hepatic *Pck1* and *G6pc* by 6- and 2-fold (Figure 2C), respectively, and PCK1 and G6PC protein levels 2-fold (Figure 2D–E). Consistent with this, knockdown of hepatic TAZ significantly increased the *ad libitum*-fed and fasting blood glucose concentrations (Figure 2F). Mice with hepatic TAZ knockdown also showed larger blood glucose excursions than control mice when challenged with pyruvate, a gluconeogenic substrate (Figure 2G). By contrast, the knockdown of TAZ did not alter the levels of YAP or key gluconeogenic factors (e.g., hepatic nuclear receptor alpha (HNF4α), forkhead box O1 (FoxO1), and GR)^29^, suggesting that the inhibition of gluconeogenic gene by TAZ is unlikely to occur *via* regulation of the protein levels of these key transcription factors (Figure 2–Figure supplement 2). It also had little effect on the phosphorylation of AKT, FoxO1, and CREB (Figure 2–Figure supplement 2), or on the plasma concentrations of insulin or glucagon (Figure 2–Figure supplement 3), hormones that regulate gluconeogenic gene expression.

**Figure 2.**
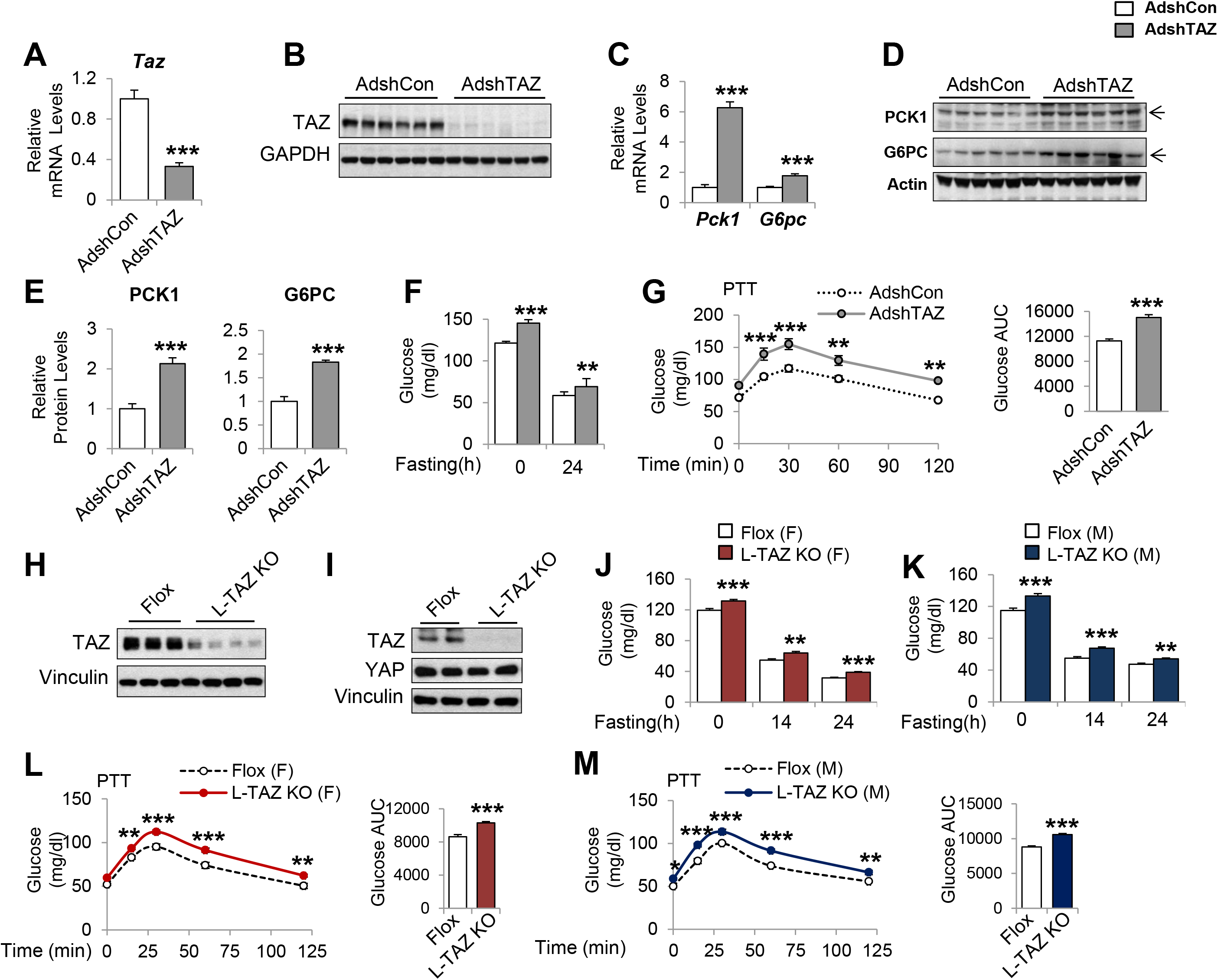
Knockdown or knockout of hepatic TAZ increases gluconeogenic gene expression and blood glucose concentrations in mice. (A–F) Eight- to twelve-week-old C57BL/6J mice were administered AdshTAZ or AdshControl (AdshCon), then sacrificed 8 days later in the *ad libitum*-fed state, when the mRNA expression of hepatic genes (A and C) and the corresponding protein levels (B, D, and E [quantified results of D]) were measured. (F) *Ad libitum*-fed and fasting blood glucose concentrations were measured. (G) Six days after adenoviral injection, the mice underwent pyruvate tolerance testing (PTT) (left) and the areas under the curves (AUC) were calculated (right). (H–M) Ten- to twelve-week-old, age- and sex-matched L-TAZ KO and floxed (Flox) controls were studied. (H–I) TAZ protein was measured in whole liver cell lysates (H) or isolated hepatocytes (I) by immunoblotting. (J–K) *Ad libitum*-fed and fasting blood glucose concentrations were measured. (L–M) Mice underwent PTT. Data are means and SEMs; control values were set to 1; n = 7–9. Representative results of two to three independent experiments are shown. Data were analyzed by unpaired Student’s *t*-test (A, C, E–F, G (right), J–K, and L–M (right)) and two-way ANOVA (G and L–M (left)); \**p* < 0.05, \*\**p* < 0.01, and \*\*\**p* < 0.001.

To validate these results, we used a genetic TAZ deletion model, L-TAZ KO mice (TAZ^flox/flox^:Albumin-Cre), in which TAZ is deleted in fetal liver. L-TAZ KO mice were born at Mendelian ratios and showed little difference in body mass from sex and age-matched littermate floxed controls (Flox) (Figure 2–Figure supplement 4A). The knockout of TAZ was confirmed by immunoblotting lysates prepared from liver and isolated hepatocytes (Figure 2H–I). The knockout of TAZ also had no overt effects on liver mass or histology (Figure 2–Figure supplement 4B–C). Previous studies showed that TAZ deletion in phosphatase tension and homolog (PTEN) knockout mice reduces insulin receptor substrate (IRS) 2 expression in mouse liver^30^ and that TAZ deletion in muscle reduces IRS1 expression^31^. In addition, it has been shown that TAZ deletion in white adipose tissue improves insulin sensitivity^32^. However, L-TAZ KO mice did not have differing levels of IRS1 and 2 expression or insulin sensitivity relative to floxed controls (Figure 2–Figure supplement 5A–B). Nonetheless, similar to the aforementioned AdshTAZ-treated mice, both male and female L-TAZ KO mice had higher *ad libitum*-fed and fasting blood glucose concentrations (Figure 2J–K). These knockout mice also showed a larger increase in blood glucose during pyruvate tolerance testing (PTT) (Figure 2L–M). Taken together, these data show that deletion of hepatic TAZ produces a fasting-like state in the liver, in which gluconeogenic gene expression and glucose production are induced.

### Hepatic overexpression of TAZ reduces gluconeogenic gene expression and blood glucose in mice

To determine whether overexpression of TAZ is sufficient to inhibit the expression of gluconeogenic genes in the fasting state, we constructed an adenovirus expressing FLAG epitope-tagged human TAZ (AdTAZ), which expresses a slightly larger protein than endogenous mouse TAZ (Figure 3A–B). Immunohistochemistry confirmed the expression of AdTAZ in both pericentral and periportal hepatocytes (Figure 3–Figure supplement 1) and immunoblotting revealed that the level of exogenously expressed TAZ was comparable to endogenously expressed TAZ in pericentral hepatocytes (Figure 3B). Compared with the administration of control virus expressing green fluorescent protein (AdGFP), AdTAZ administration caused relative hypoglycemia during fasting (Figure 3C), which was accompanied by lower mRNA expression and protein levels of hepatic PCK1 and G6PC (Figure 3D–E). TAZ overexpression did not affect mouse body or white adipose mass, but modestly increased liver mass, without affecting liver histology (Figure 3–Figure supplement 2A–D). TAZ overexpression also blunted the rise in blood glucose caused by the injection of glucagon or pyruvate (Figure 3G–H), without affecting the protein levels of key gluconeogenic factors, the phosphorylation of AKT or CREB, or plasma insulin or glucagon concentrations (Figure 3–Figure supplement 2 E–F). Taken together, these data indicate that the overexpression of TAZ in the fasting state mimics the fed state and is sufficient to inhibit hepatic gluconeogenic gene expression and glucose production.

**Figure 3.**
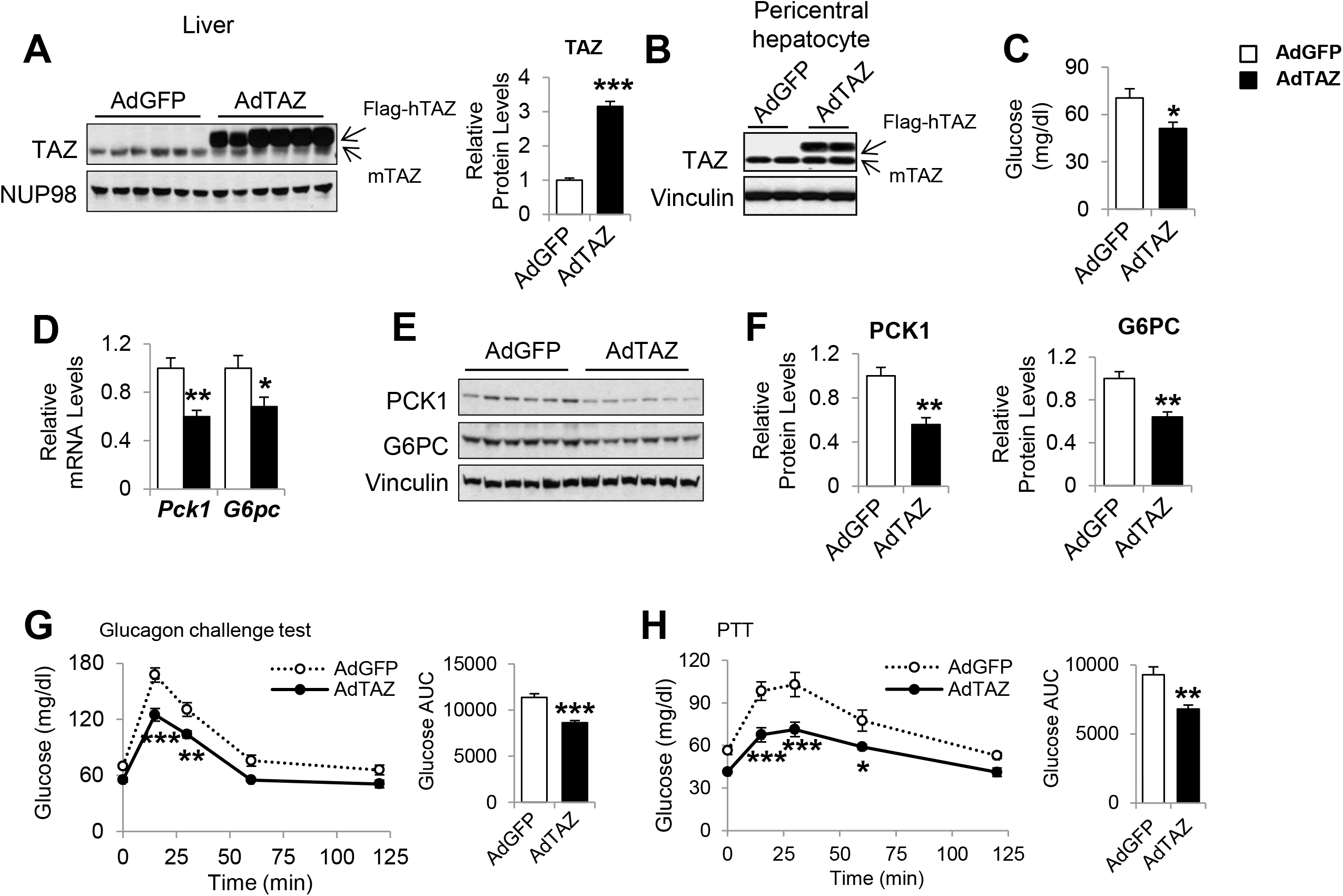
Overexpression of hepatic TAZ inhibits gluconeogenic gene expression and reduces blood glucose concentration in mice. Eight- to twelve-week-old C57BL/6J mice were administered AdGFP or AdTAZ, then sacrificed 5 days later, after a 24-h fast, when the hepatic protein levels in nuclear extracts (A [representative image, left; quantified results, right]), isolated pericentral hepatocytes (B), or whole cell lysates (E and F [quantified results of E]) were measured. Blood glucose (C), and mRNA expression of hepatic genes (D) were measured. (G–H) Five days after adenoviral injection, the mice underwent glucagon challenge (G) or PTT (H). Data are means and SEMs; control values were set to 1; n = 5–10. Representative results of two to three independent experiments are shown. Data were analyzed by unpaired Student’s *t*-test (A, C–D, F, and G–H (right) and two-way ANOVA (G–H (left)); \**p* < 0.05, \*\**p* < 0.01, and \*\*\**p* < 0.001.

### The regulation of gluconeogenic gene expression by TAZ is hepatocyte-autonomous

Infection of mouse primary hepatocytes with AdshTAZ reduced both TAZ mRNA expression and protein level (Figure 4A), and markedly increased the glucagon and dexamethasone (Dex)-induced mRNA expression of *Pck1* and *G6pc* (Figure 4B–C). TAZ knockdown also increased glucose secretion into the medium by glucagon-stimulated hepatocytes (Figure 4D), which demonstrates the functional importance of these changes in gene expression. Conversely, the infection of mouse primary hepatocytes with AdTAZ increased TAZ protein levels 3-fold compared with infection with control AdGFP (Figure 4E), and substantially inhibited the glucagon and dexamethasone-induced mRNA expression of *Pck1* and *G6pc* (Figure 4F–G). Consistent with these data, AdTAZ infection also reduced glucose production after glucagon stimulation (Figure 4H). In addition, knockdown or overexpression of TAZ did not alter the protein levels of YAP or gluconeogenic transcription factors (Figure 4–Figure supplement 1). Given that insulin and glucagon are major regulators of gluconeogenic genes, we determined whether TAZ regulates their action in hepatocytes, and found that it had no effects on either the basal or stimulated phosphorylation of CREB or AKT in hepatocytes treated with glucagon or insulin, respectively (Figure 4–Figure supplement 2), suggesting that TAZ is unlikely to directly affect gluconeogenic gene expression by altering the cellular sensitivity to these hormones in hepatocytes.

**Figure 4.**
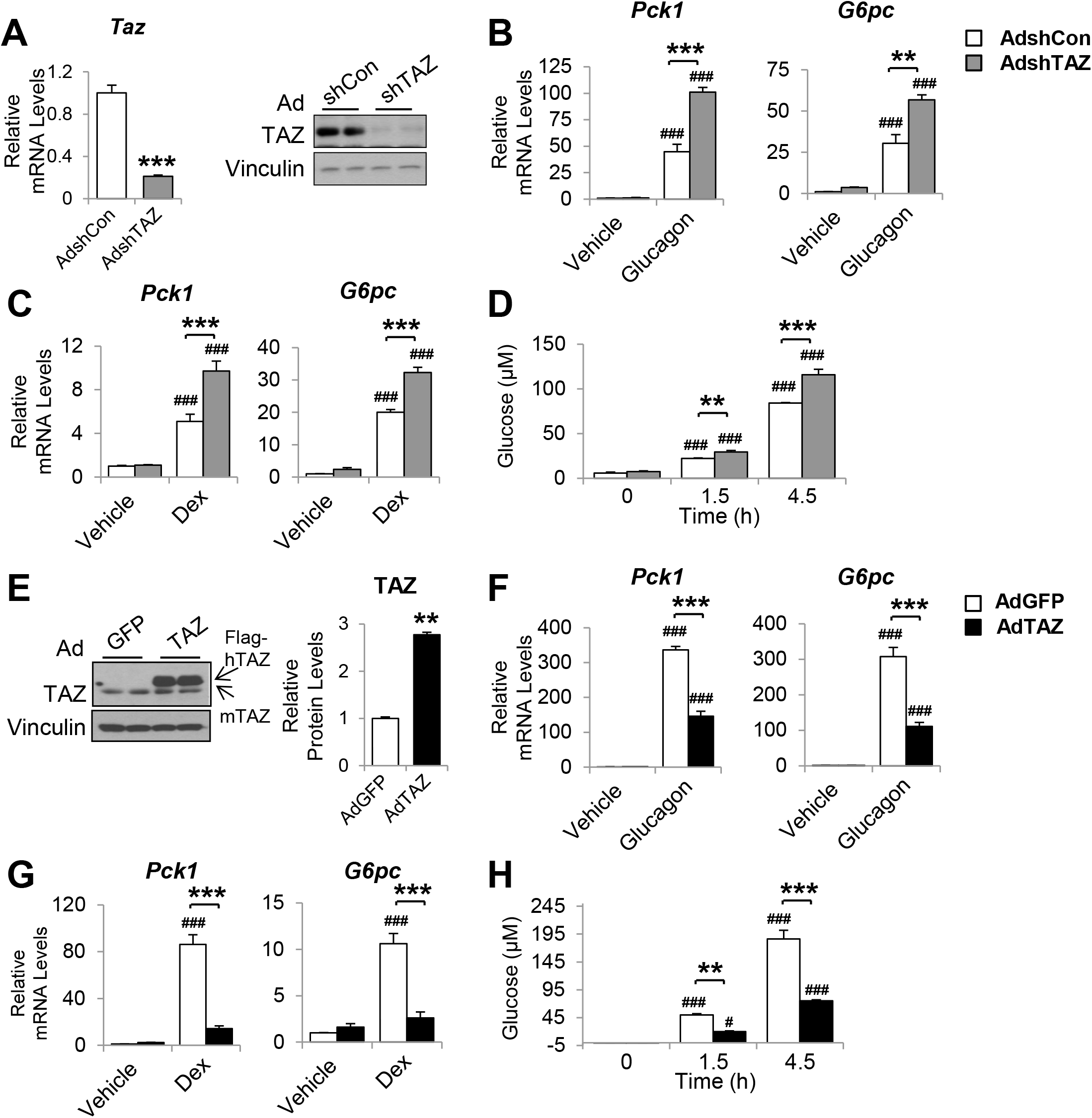
The inhibition of gluconeogenic gene expression by TAZ is hepatocyte-autonomous. Primary mouse hepatocytes were isolated from eight- to twelve-week-old C57BL/6J mice. (A–D) To knockdown TAZ, cells were infected with AdshTAZ or AdshControl (AdshCon). (E–H) To overexpress TAZ, cells were infected with AdTAZ or AdGFP. Cells were treated with glucagon (20 nM) for 3 h (B, F) or dexamethasone (Dex, 100 nM) for 6 h (C, G). (A (left)–C, F, and G) Gene expression was measured using real-time RT-PCR. (A (right) and E) TAZ protein levels were measured by immunoblotting whole cell lysates and the quantified results of E are shown on the right. (D and H) Glucose production, in the presence of glucagon (20 nM), was assessed by measuring the glucose concentration in the media at the indicated times. Data are means and SEMs of three wells; control values were set to 1. Data were analyzed by unpaired Student’s *t*-test (A and E) and two-way ANOVA (B–D and F–H). \**p* < 0.05, \*\**p* < 0.01, and \*\*\**p* < 0.001 *versus* control adenovirus-treated cells; in B–D and F–H, #*p* < 0.05 and ###*p* < 0.001 *versus* similarly-treated controls. Representative results of two to five independent experiments are shown.

### TAZ inhibits the GR-transactivation of gluconeogenic genes

We next determined the molecular mechanisms by which TAZ inhibits gluconeogenic gene expression. Because TAZ is not known to bind directly to gluconeogenic gene promoters, we determined whether the transcriptional effects of TAZ are dependent on other factors in cell culture. Consistent with the results of previous studies, co-transfection of HepG2 cells with vectors expressing GR, HNF4α, or PGC1α, followed by Dex treatment, induced human *G6PC* and *PCK1* luciferase reporter activity (*G6PC*-Luc and *PCK1*-Luc, respectively) (Figure 5A and Figure 5–Figure supplement 1A). Overexpression of TAZ in these cells inhibited the activity of the *G6PC* and *PCK1* promoters by 50% (Figure 5A and Figure 5–Figure supplement 1A), indicating that TAZ suppresses transcription of these genes. To determine whether the Hippo pathway and TEADs are required for these effects of TAZ, we utilized TAZS89A^25^ and TAZS51A^33^ mutants, which abolish TAZ phosphorylation by LATS1/2 and cannot interact with TEADs, respectively. Both mutants inhibited *G6PC-* Luc to a similar extent, compared to wild-type TAZ (Figure 5A), suggesting that the Hippo pathway and TEADs are not required. Conversely, the transfection of cells with a vector expressing an shRNA targeting human TAZ reduced TAZ expression (Figure 5–Figure supplement 1B) and increased the activity of *G6PC*-Luc and *PCK1*-Luc by >40% (Figure 5B and Figure 5–Figure supplement 1C).

**Figure 5.**
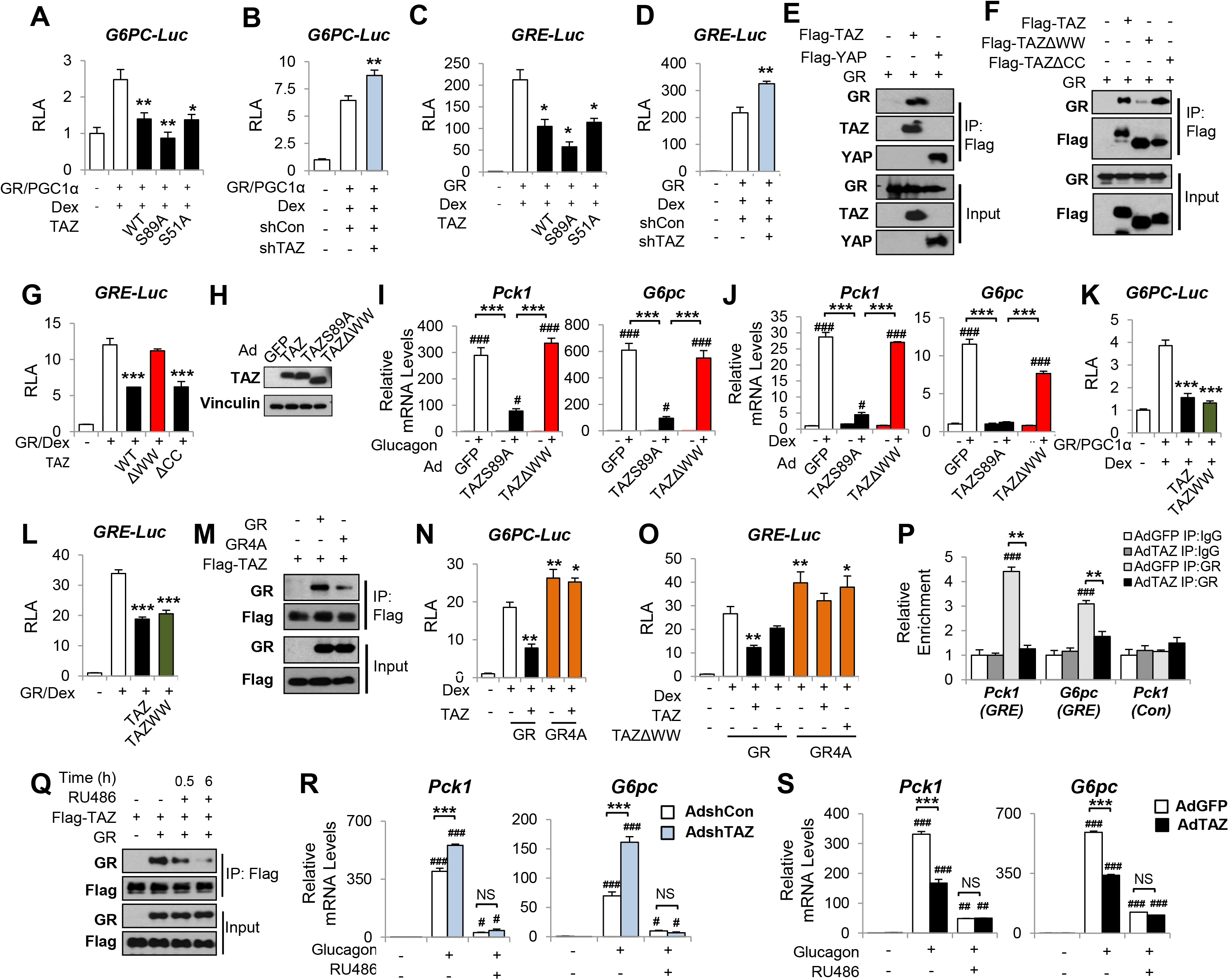
TAZ interacts with GR and inhibits the transactivation of gluconeogenic genes by GR. (A–D, G, K–L, and N–O) HepG2 cells were co-transfected with expression vectors, luciferase reporters, and an internal control (*Renilla*), and treated with Dex (100 nM), as indicated. Relative Luciferase Activity (RLA) is presented after normalization to the *Renilla* activity. (E–F, M, and Q). 293A Cells were transfected with expression vectors, which was followed by immunoprecipitation and immunoblotting, as indicated. (H–J and R–S) Primary mouse hepatocytes were infected with adenoviruses, with or without glucagon or Dex treatment. (R–S) Cells were treated with RU486 30 min prior to glucagon treatment. Protein levels (H) and gene expression (I–J and R–S) were measured by immunoblotting and real-time RT-PCR, respectively. (P) ChIP assays were performed using an anti-GR antibody or a control IgG. The relative enrichment of GR was assessed using real-time RT-PCR and primers for the indicated regions of the *Pck1* and *G6pc* genes. Data are means and SEMs of three or four wells or immunoprecipitation reactions. Representative results of two to five independent experiments are shown. Data were analyzed by one-way ANOVA (A–D, G, K–L, and N–O) and two-way ANOVA (I–J, P, and R–S). \**p* < 0.05, \*\**p* < 0.01, and \*\*\**p* < 0.001; #*p* < 0.05, ##*p* < 0.01, and ###*p* <0.001. In A–D, G, K–L, and N–O, * denotes comparisons with wells treated with GR/Dex or GR/PGC1α/Dex alone; in I–J and R–S, # denotes comparisons with controls administered the same viruses; in P, # denotes a comparison with IgG.

To identify the molecular target of TAZ, we constructed a luciferase reporter containing three repeats of a canonical GR response element (inverted hexameric half-site motifs, separated by a three-base-pair spacer) (*GRE*-Luc)^34^. Similar to our findings using *G6PC*-Luc and *PCK1*-Luc, TAZ suppressed dexamethasone-stimulated *GRE*-Luc activity by >50%, whereas knockdown of TAZ increased promoter activity by 40% (Figure 5C–D), suggesting that TAZ inhibits GR transactivation.

### The interaction of TAZ with GR is dependent on its WW domain

The WW domain of TAZ has been reported to interact with proteins containing an I/L/PPxY (I, isoleucine; L, leucine; P, proline; x, any amino acid; and Y, tyrosine) motif^35^ ^21^. Moreover, an I/LPxY motif found in the ligand binding domain of GR is highly conversed across species, including in humans, mice, and rats (Figure 5–Figure supplement 2). Co-immunoprecipitation (co-IP) assay revealed that TAZ, but not YAP, was able to interact with GR (Figure 5E). These data are consistent with our previous findings that YAP does not interact with PGC1α^23^, a co-activator of GR. A TAZ mutant protein lacking the WW domain (TAZΔWW), but not one lacking the coiled-coil (CC) domain (TAZΔCC), showed much weaker interaction with GR (Figure 5F) and was unable to inhibit *GRE*-Luc (Figure 5G). Consistent with this, compared with the TAZS89A mutant that suppressed glucagon or Dex-induced activation of *Pck1* and *G6pc* gene expression by >80% in primary mouse hepatocytes, TAZΔWW failed to suppress glucagon or Dex-induced *Pck1* and *G6pc* expression (Figure 5H–J). Furthermore, a TAZ mutant consisting of only the WW domain (TAZWW) was sufficient to inhibit *G6PC*-Luc and *GRE*-Luc activity (Figure 5K–L), but was unable to activate TEAD, due to the lack of a TEAD-binding domain (Figure 5–Figure supplement 3), suggesting that the effects of TAZ on gluconeogenic genes can be separated from its effects on proliferative genes.

Conversely, a GR mutant in which the conserved IPKY motif was mutated to alanines (A) (GR4A mutant) interacted with TAZ much more weakly, despite being expressed at a level similar to that of the wild-type GR (Figure 5M). In addition, compared with wild-type GR, the GR4A mutant was not subject to TAZ-mediated inhibition of *GRE*-Luc and displayed a greater ability to induce the activities of the *GRE*-Luc and *G6PC*-Luc reporters (Figure 5N–O).

### TAZ inhibits the binding of GR to GREs

To understand how the interaction between TAZ and GR inhibits GR transactivation, we determined how the binding of TAZ to GR inhibits GR nuclear localization, dimerization, and binding to promoter GREs. Whereas Dex treatment induced the nuclear accumulation of GR, overexpression of TAZ had little effect on its subcellular distribution in either the absence or presence of Dex (Figure 5–Figure supplement 4A). Similarly, TAZ did not reduce the quantity of dimeric GR in cells, as shown by immunoblotting after the treatment of cells with a cross-linker (dithiobis (succinimidyl propionate), DSP) (Figure 5–Figure supplement 4B). In addition, the fact that the GR4A mutant can be activated by Dex (Figure 5N–O) also suggests that the TAZ-GR interaction does not impair GR nuclear localization, ligand activation, or dimerization.

To evaluate GR binding to the *G6pc* and *Pck1* promoters, we performed chromatin immunoprecipitation (ChIP) assays in primary mouse hepatocytes using control IgG and anti-GR antibodies. Glucagon treatment strongly promoted the binding of GR to the *Pck1* and *G6pc* promoter regions containing GREs (‘*Pck1* (GRE)’, −376 to −280 and ‘*G6pc* (GRE)’, −215 to −111), but not to a distal region of the *Pck1* promoter lacking GREs (‘*Pck1* (Con)’, −3,678 to −3,564). However, the glucagon-induced binding of GR to the *Pck1* or *G6pc* promoters was significantly lower in cells overexpressing TAZ (Figure 5P). Taken together, these data suggest that the TAZ-GR interaction limits gluconeogenic gene expression by reducing GR binding to GREs.

To determine whether TAZ might have significant GR-independent effects to reduce hepatic gluconeogenic gene expression, we treated primary mouse hepatocytes with RU486, a well-characterized and potent GR antagonist^36^. Upon binding to GR, RU486 induces a conformational change that favors the interaction of GR with co-repressors, but stabilizes the binding of dimeric GR to GREs^36^. Interestingly, we found that RU486 blocked the interaction of GR with TAZ (Figure 5Q), probably due to the change in GR conformation. As expected, RU486 treatment of hepatocytes reduced glucagon-stimulated *Pck1* and *G6pc* mRNA expression (Figure 5R–S), whereas the knockdown of TAZ increased glucagon-stimulated *Pck1* and *G6pc* expression in the absence of RU486. However, this increase was entirely abolished in the presence of RU486 (Figure 5R). Similarly, RU486 prevented the inhibitory effects of TAZ overexpression on *Pck1* and *G6pc* expression in hepatocytes (Figure 5S). Thus, not only is the TAZ-GR interaction necessary for the reduction in gluconeogenic gene expression, but it is likely the sole mechanism for such a reduction.

### TAZ inhibits GR transactivation in mice

Consistent with the interaction between GR and TAZ identified in cultured cells, GR in the nuclear extracts of wild-type mouse liver could be immunoprecipitated using an antibody targeting endogenous TAZ (Figure 6A). To verify that TAZ regulates the association of GR with promoter GREs in mouse liver, we conducted ChIP assays of endogenous gluconeogenic gene promoters. GR bound to regions of the *G6pc* and *Pck1* promoters containing functional GREs (Figure 6B). The binding of GR to GRE-containing regions of the *G6pc* and *Pck1* promoters was significantly increased by TAZ knockdown (Figure 6B). Moreover, the histone acetylation of *G6pc* and *Pck1* promoter regions containing or near to GREs was increased 50–80% by TAZ knockdown (Figure 6C), implying active transcription from the *G6pc* and *Pck1* genes. Conversely, ChIP assays revealed that TAZ overexpression caused a 25–45% reduction in the binding of GR to GREs, histone acetylation, and the binding of the polymerase II subunit (Pol II) to the promoter regions of *G6pc* and *Pck1* (Figure 6D–F). However, TAZ bound normally to the promoter of *Ctgf* (a well-described TAZ target gene that is not known to be regulated by GR) in AdTAZ-infected mice, but not to the promoters of *G6pc* or *Pck1* (Figure 6– Figure supplement 1). These data support a model in which TAZ inhibits GR transactivation by interacting with GR and suppressing GR binding to GREs, rather than a model in which TAZ binding near or within GREs inhibits the binding of GR to gluconeogenic gene promoters.

**Figure 6.**
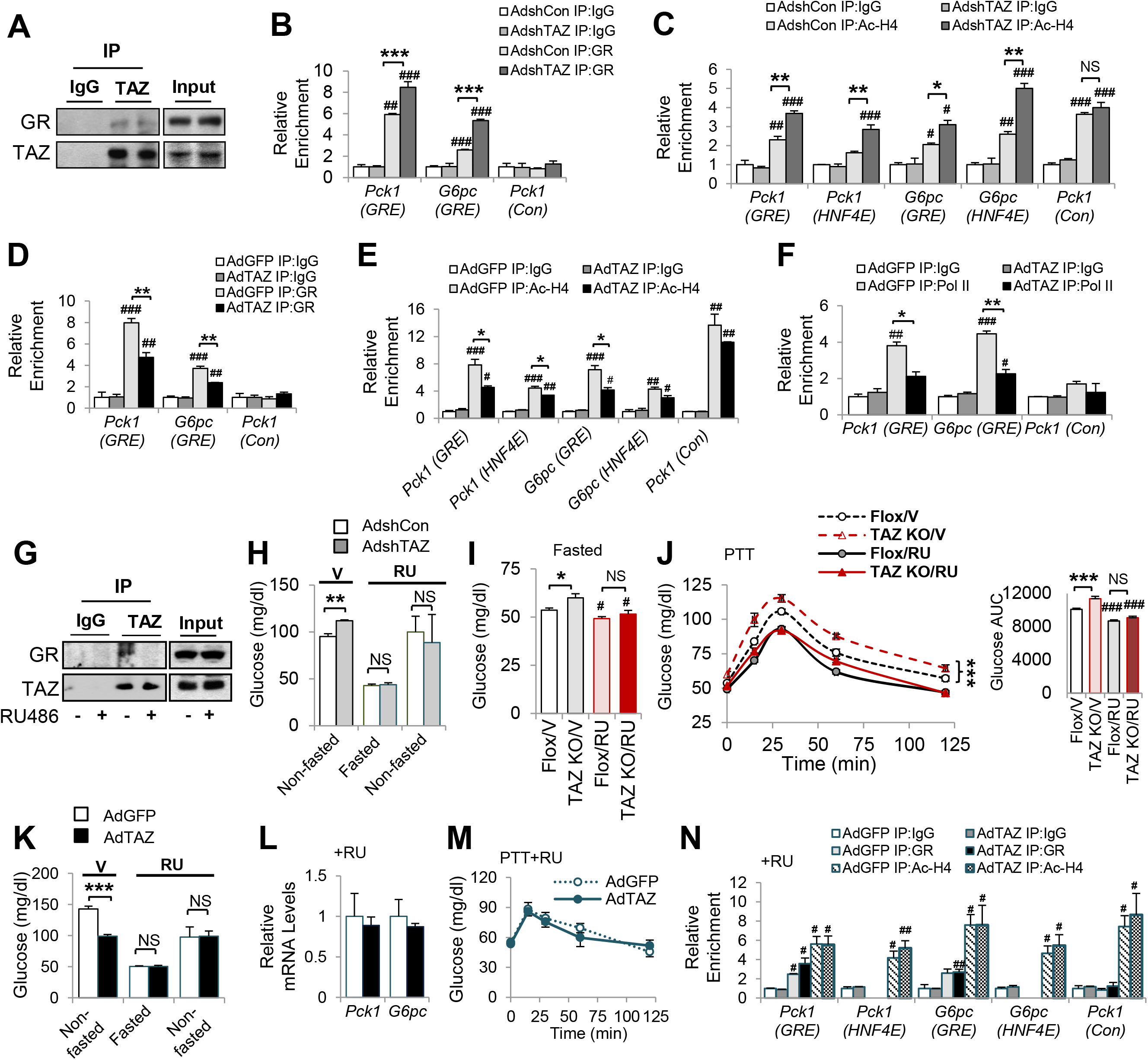
TAZ represses GR-transactivation of gluconeogenic genes in mouse liver. (A and G) Endogenous TAZ was immunoprecipitated from liver nuclear extracts prepared from *ad libitum*-fed C57BL/6J mice (A) or mice treated with RU486 or vehicle (G), and the amounts of GR in the immunoprecipitates were measured by immunoblotting. C57BL/6J mice administered AdshTAZ or AdshCon, as in Figure 2 (B–C, and H) or AdTAZ or AdGFP, as in Figure 3 (D–F and K–N). (I–J) L-TAZ KO and control mice. (H–N) Mice were treated with RU486 (RU) or vehicle (V), as indicated. The relative enrichment of GR (B, D, and N), acetylated-histone 4 (Ac-H4) (C, E, and N), and Pol II (F) in the liver extracts was assessed using ChIP assays. (H, I, and K) Blood glucose concentration. (L) Hepatic gene expression. (J and M) PTT. Data are means and SEMs; n=5–8; for ChIP, the data are the results of triplicate or quadruplicate immunoprecipitations. Data were analyzed by two-way ANOVA (B–F, I–J and M–N) and unpaired Student’s *t*-test (H and K–L). \**p* < 0.05, \*\**p* < 0.01, and \*\*\**p* < 0.001; #*p* < 0.05, ##*p* < 0.01, and ###*p* < 0.001; in B–F and N, # denotes comparisons with IgG; in I and J, # denotes comparisons with vehicle-treated controls of the same genotype; NS, not significant.

We next determined whether the inhibition of gluconeogenic gene expression by TAZ requires GR. Because RU486 reduced the TAZ-GR interaction in cultured cells (Figure 5Q) and mouse liver nuclear extracts (Figure 6G), we determined whether RU486 could abrogate the effects of TAZ on glucose homeostasis in mice. As expected, TAZ knockdown increased mouse blood glucose concentration prior to RU486 injection. However, this increase was completely abolished by RU486 treatment (Figure 6H). Similarly, hepatic TAZ knockout increased fasting blood glucose and glucose production after pyruvate administration to mice; and RU486 treatment abolished the effects of TAZ on blood glucose and glucose production (Figure 6I–J). Consistent with these loss-of-function studies, RU486 also entirely abolished the ability of overexpressed hepatic TAZ to reduce gluconeogenic gene expression, to improve pyruvate tolerance, and to inhibit gluconeogenic gene expression (Figure 6K–M). Moreover, RU486 abolished the ability of TAZ to reduce GR binding to GREs in the *Pck1* or *G6pc* promoters and the amounts of acetylated histones on these promoters (Figure 6N). These data confirm that the interaction between GR and TAZ is required for the inhibitory effects of TAZ on hepatic gluconeogenic gene expression in mice.

The binding of dimeric GR to GREs is required for the GR transactivation of gluconeogenic genes, but not GR transrepression of anti-inflammatory genes ^15^. Therefore, we next determined the effects of TAZ on the regulation of gluconeogenic and inflammatory genes by Dex in mouse liver. TAZ overexpression inhibited the Dex-induced increases in blood glucose and gluconeogenic gene expression, but the effects of TAZ on the inhibition of inflammatory gene expression by Dex, including the expression of the tumor necrosis factor alpha (*Tnfα*) and interleukin 1 (*Il1*) genes, were inconsistent (Figure 7–Figure supplement 1). This inconsistency may be the result of the ability of TAZ to directly induce hepatic inflammation^37^.

To determine whether the TAZ WW domain is required for GR regulation *in vivo*, we expressed TAZΔWW, TAZS89A, or GFP in mouse liver. AdTAZΔWW administration had no effects on mouse body and liver mass (Figure 7–Figure supplement 2A–B). Compared with control AdGFP, TAZS89A reduced the blood glucose of *ad libitum*-fed, fasted, or pyruvate-challenged mice (Figure 7A–B). By contrast, TAZΔWW failed to reduce blood glucose and glucose production during PTT (Figure 7A–B), although the TAZΔWW mutant appeared to be more stable in mouse liver (Figure 7–Figure supplement 2C). In addition, the TAZΔWW mutant failed to inhibit *Pck1* and *G6pc* expression and the binding of GR to GREs in the *Pck1* and *G6pc* promoters in mouse liver (Figure 7C–D). Taken together, these data suggest that the WW domain mediates the inhibition of gluconeogenic gene expression by TAZ *in vivo*.

**Figure 7.**
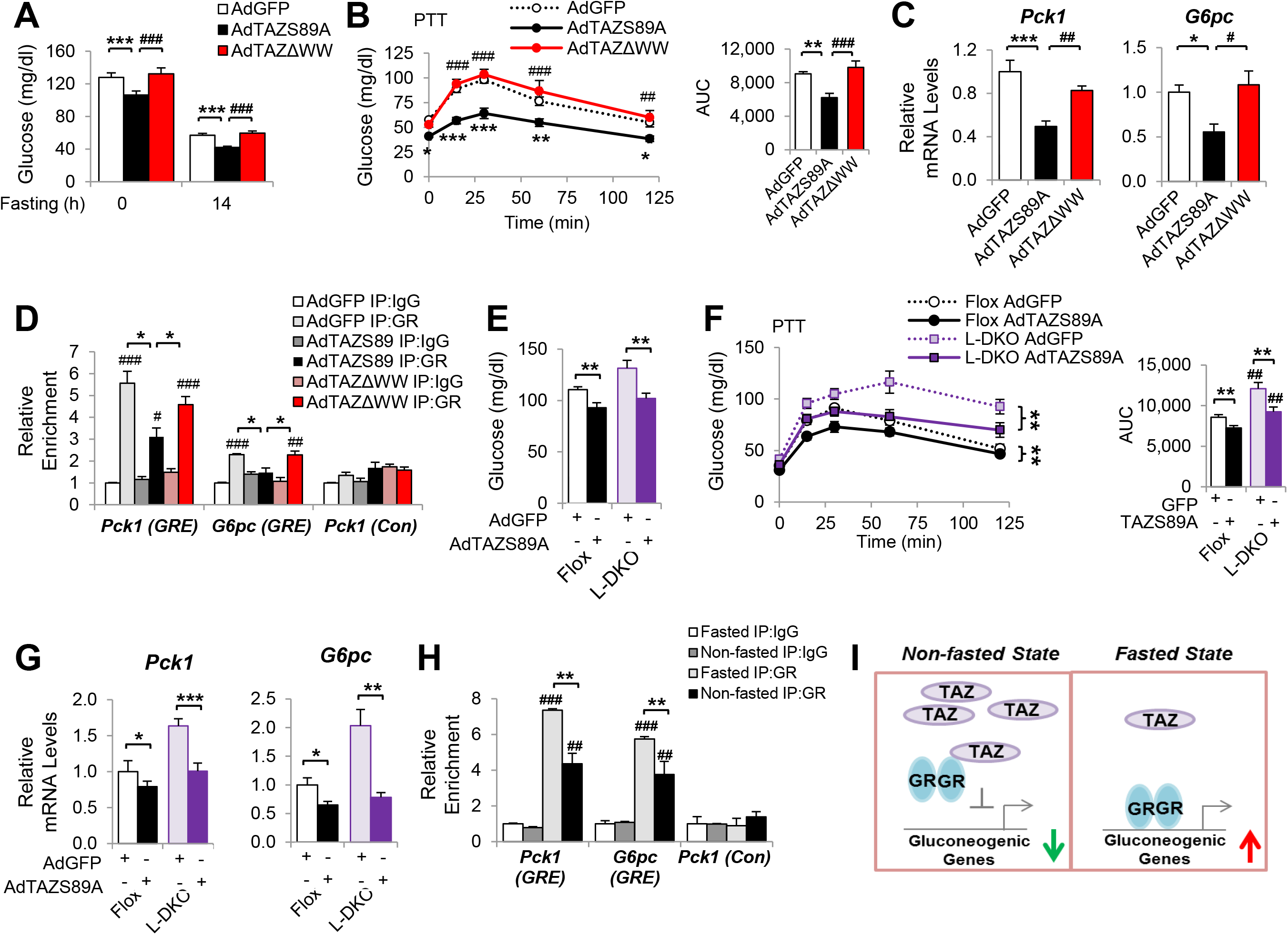
The inhibition of gluconeogenic gene expression by TAZ requires its WW domain and is independent of insulin signaling in mice. C57BL/6J mice (A–D) or L-DKO or flox controls (E–G) were administered the indicated adenoviruses. (H) C57BL/6J mice were *ad libitum*-fed (Non-fasted) or fasted for 24 h. (A and E) Blood glucose. (B and F) PTT. (C and G) Hepatic gene expression, (D and H) ChIP assays were conducted using the indicated antibodies. (I) Regulation of gluconeogenic gene expression by TAZ in the fed and fasting states. Data are means and SEMs; n = 5–8; except for D and H, in which the data are the results of triplicate or quadruplicate immunoprecipitations. Data were analyzed by one-way ANOVA (A, B (right), and C), two-way ANOVA (B (left), D, F, and H), and unpaired Student’s *t*-test (E and G). \**p* < 0.05, \*\**p* < 0.01, and \*\*\**p* < 0.001; #*p* < 0.05, ##*p* < 0.01, and ###*p* < 0.001. In B, * denotes comparisons with AdGFP and # denotes comparisons between AdS89A and AdTAZΔWW; in D and H, # denotes comparisons with IgG; in F, # denotes comparisons with flox mice administered the same virus.

TAZ has previously been shown to regulate IRS2 expression in liver cancer^30^ and to alter insulin sensitivity^32^, but we did not observe effects of TAZ on IRS1/2 expression or insulin sensitivity in mice (Figure 2–Figure supplement 5). However, we determined whether the inhibition of gluconeogenic gene expression by TAZ requires insulin signaling, using liver-specific IRS 1 and 2 double knockout (L-DKO) mice. AdTAZS89A administration affected mouse body and liver mass to a similar extent in both L-DKO and floxed controls (Figure 7–Figure supplement 3A–B). AdTAZS89A also reduced blood glucose concentration, glucose production, and gluconeogenic gene expression in both L-DKO and floxed control mice (Figure 7E–G). Taken together, these data suggest that the regulation of gluconeogenic genes by TAZ does not require insulin signaling.

Fasting increases GR binding to gluconeogenic gene promoters in mouse liver and feeding reduces this binding^38^. Consistent with this, we found that the binding of GR to GREs in the *G6pc* and *Pck1* promoters was reduced by 40% in the livers of mice that were fasted, *versus* those that were fed (Figure 7H). Collectively, our data suggest a role for hepatic TAZ in glucose homeostasis in normal mouse liver. In the fed state, high hepatic TAZ expression inhibits GR transactivation of gluconeogenic genes by interacting with GR and reducing the binding of GR to the promoters of these genes; whereas in the fasted state, lower TAZ expression enables GR to bind to gluconeogenic genes and activate their transcription, which increases glucose production (Figure 7I).

## Discussion

Distinct cellular functions of YAP and TAZ have been suggested by several studies^17, 18, 39, 40^. In the liver, YAP is expressed at low levels in normal hepatocytes, which is crucial to the maintenance of their differentiated state^24^. Consistent with this, the knockout of hepatic YAP in adult mouse liver has no effect on gluconeogenic gene expression or glucose homeostasis^23^. However, hepatic YAP expression negatively correlates with the expression of gluconeogenic genes in human hepatocellular carcinoma, which suggests a role for YAP in the integration of glucose metabolic regulation and cell growth^23^. By contrast, TAZ is expressed in normal pericentral hepatocytes and regulates normal glucose homeostasis. The WW domain of TAZ is required for its interaction with GR, because a TAZ mutant lacking this domain cannot interact with GR and does not inhibit the transactivation of gluconeogenic genes by GR. Interestingly, although YAP also contains WW domains, it does not interact with GR (Figure 5E) or a GR co-activator, PGC1α^23^. Therefore, these results demonstrate the distinct roles of TAZ and YAP in the modulation of GR activity and hepatic physiology.

The essential roles of GR and GCs in the promotion of hepatic gluconeogenic gene transcription are well established. Hepatocyte-specific GR ablation leads to the death of half of newborn mice and severe hypoglycemia in adult mice because of a defect in gluconeogenesis^7^. In addition, GR is required for the full induction of glucose production, because fasting, glucagon, cyclic AMP (cAMP), and epinephrine-activated glucose production are substantially blunted in adrenalectomized rodents^8, 9^. The binding of GR to the promoters of gluconeogenic genes is higher in the fasting state than in the fed state^38^, and GCs are believed to play a role in this dynamic regulation. However, little is known about the GC-independent factor(s) or mechanism(s) mediating this regulation. Our data reveal hepatic TAZ to be a novel regulator of GR that modulates the binding of GR to gluconeogenic gene promoters. First, TAZ inhibits GR transactivation. The overexpression of TAZ inhibits, while the knockdown of TAZ increases, the promoter activity of *GRE*-Luc, *G6PC*-Luc, and *PCK1*-Luc reporter genes. In addition, TAZ inhibits the induction of gluconeogenic gene expression by Dex. Second, TAZ acts as a repressor of GR. However, unlike a classical co-repressor of GR, such as nuclear receptor compressor 1 (NCoR1), which does not alter DNA-binding by GR^41^, the binding of TAZ to GR results in the dissociation of GR itself from the promoters of GR target genes. Consistent with this, the overexpression of TAZ reduces GR binding to GREs in the promoters of gluconeogenic genes, whereas the knockdown of TAZ increases GR binding to these GREs. Third, the effects of TAZ overexpression or knockdown on gluconeogenic gene expression and the binding of GR to the *Pck1* and *G6pc* promoters are completely abolished in hepatocytes in the presence of RU486, which prevents the interaction of GR with TAZ.

RU486, a GR antagonist, binds to GR but elicits conformational changes that do not favor the recruitment of histone acetyltransferase without an inhibition of GR nuclear localization and DNA-binding^36^. Thus, one explanation for the reduction in the interaction between GR and TAZ following RU486 treatment is that RU486-bound GR possesses a conformation that does not permit GR to interact with TAZ. The ligand binding domain of GR is crucial for the interaction of GR with co-regulators and ligand-binding, but also for the formation of homodimers, which permit GR to bind to GREs in gluconeogenic gene promoters. Our results suggest that the binding of TAZ to the LBD of GR does not lead to a reduction in the nuclear localization of GR or in its ability to respond to an agonist; however, exactly how the binding of TAZ to GR causes GR to dissociate from GRE is unknown. GRβ, which is produced by alternative splicing, differs only at its C-terminus from GRα, where the I/LPxY motif resides. Interestingly, unlike GRα, GRβ is not able to bind to conventional GREs in gluconeogenic gene promoters and activate gluconeogenic gene transcription^42^, which implies that these C-terminal amino acids are indispensable for the binding of GR to GREs.

The inhibition of gluconeogenic gene expression by TAZ is hepatocyte-autonomous and is unlikely to be an indirect effect of glucagon or insulin signaling. TAZ does not affect the phosphorylation of CREB in hepatocytes, nor are plasma glucagon concentrations altered when TAZ is overexpressed or knocked down. Previous studies show that the overexpression of TAZ increases insulin signaling in liver cancer and that TAZ deletion in white adipose and muscle increases insulin sensitivity^25, 30^. However, we found little effect of either TAZ overexpression or knockdown on the phosphorylation of AKT or FoxO1 (Ser256), or on the plasma insulin concentration and we found little effects of hepatic TAZ knockout on whole-body insulin sensitivity. In addition, the overexpression of hepatic TAZ is sufficient to reduce blood glucose, pyruvate tolerance, and the hepatic expression of *Pck1* and *G6pc* in liver-specific IRS 1 and 2 double-knockout mice^43^, which strongly suggests that the inhibition of the GR transactivation of gluconeogenic genes by TAZ is independent of hepatic insulin signaling.

TAZ protein, but not mRNA expression, was altered by fasting and feeding, which suggests that TAZ is subject to post-transcriptional regulation. It is also noteworthy that TAZ is zonally distributed, with the highest expression being near the central veins and the lowest near the hepatic veins. This pattern of distribution is negatively related to the expression and activity of gluconeogenic genes^26^, which is consistent with an inhibitory effect of TAZ on hepatic gluconeogenesis under normal physiologic conditions. However, the molecular mechanism(s) that mediate the post-transcriptional effects and zonal distribution of TAZ are unclear and are currently under investigation in our laboratory.

Gluconeogenesis is substantially upregulated in diabetic patients and contributes significantly to the lack of control of blood glucose concentration in these patients^4, 44^. Long-term increases in the concentration and/or actions of endogenous GCs in both humans and rodents manifest as metabolic syndrome, which is characterized by higher glucose production and insulin resistance, and resembles Cushing’s syndrome^45^. GCs are among the most widely prescribed anti-inflammatory and immunosuppressive drugs^46^. However, GC therapy is associated with substantial increases in glucose production and insulin resistance; thus, therapies that would have the anti-inflammatory effects of GCs, but not affect gluconeogenesis, would be of great interest^47^. Similarly, strategies aimed at specifically inhibiting the ability of GR to induce gluconeogenesis have been explored for the treatment of diabetes^48^. Therefore, whether the inhibition of GR by TAZ plays a role in the regulation of hepatic gluconeogenesis in these abnormal states, and whether a non-tumorigenic TAZ mutant(s), or small peptide(s) or molecule(s), which would mimic or enhance the TAZ-GR interaction could normalize the hyperglycemia associated with obesity, insulin resistance, or the chronic use of GCs, should be determined in future investigations.

In summary, the factors and mechanisms that regulate cell proliferation substantially, such as the mTOR complex^49^, overlap with those that control metabolic homeostasis, and have emerged as an important area of biology in recent years. We have identified hepatic TAZ, a downstream effector of the Hippo pathway, as a novel regulator of glucose metabolism in the normal liver. TAZ acts as a GR co-repressor and inhibits the binding of GR to GREs in the promoters of gluconeogenic genes, whereby it coordinates GR transactivation and hepatic glucose production in response to fasting and feeding, to maintain energy balance (Figure 7I).

## Acknowledgements

This project was supported by an NIH grant (DK100539) and a Boston Children’s Hospital Career Development Award (to J.M.).

## Author contributions

S.X. and J.M. conceived and designed the study. S.X., Y.L., R.H., M.W., O.S., K.H., Y.X, L. C, S.C., H.L., and L.Z. performed the experiments. All authors contributed to data analysis, and J.M. wrote the manuscript, with input from the other authors.

## Competing interests

All authors declare no conflicts of interest.

## Methods

### Animals and treatments

All mice were fed a standard chow diet and maintained on a 12 h light/dark cycle, with free access to food and water, unless otherwise indicated, and were sacrificed in the *ad libitum*-fed, fasted, or re-fed states, as indicated in the main text. At that time, the livers were removed and then fixed in 4% formaldehyde or frozen in liquid nitrogen and stored at −80°C, until RNA, protein, and immunoprecipitation analyses were performed. All animal experiments were performed with the approval of the Institutional Animal Care and Research Advisory Committee at Boston Children’s Hospital. Both male and female cohorts of the same age were studied and we found consistent results in each.

To knockdown hepatic TAZ, Eight- to twelve-week-old C57BL/6J mice were administered adenoviruses expressing shTAZ (AdshTAZ) or control shRNA (targeting human lamin) by retro-orbital injection at a dose of 1–2 × 10^9^ pfu/mouse. Six days after virus injection, mice were fasted for 14 hours and subjected to a pyruvate tolerance test (PTT), as previously described^50^. Briefly, mice were injected intraperitoneally with 2 mg/kg pyruvate at time zero, and subsequently blood glucose concentrations were measured using a glucometer (Contour). Alternatively, eight days after virus infection, the mice were sacrificed in the *ad libitum* state at zeitgeber time (ZT) 13 for gene expression and chromatin immunoprecipitation analysis. To overexpress TAZ in the liver, mice were injected with adenovirus expressing human FLAG-TAZ (AdTAZ) or control GFP (AdGFP) retro-orbitally at a dose of 0.5–1 × 10^9^ pfu/mouse. Five days later, the mice were subjected to PTT or glucagon challenge after an overnight fast or sacrificed at ZT 13 after a 24 h fast. For glucagon challenge experiment, Eight- to twelve-week-old C57BL/6J mice were injected with glucagon (250 μg/kg, Sigma) intraperitoneally at time zero, and subsequently blood glucose concentrations were measured using a glucometer. To treat RU486 (Sigma), mice were intraperitoneally injected with RU486 (50 mg/kg), fasted for 14 h, subjected to a PTT, and then sacrificed 1 day later. To treat dexamethasone (Sigma), mice were injected intraperitoneally with dexamethasone (10 μg/kg) daily for 3 days.

L-TAZ KO mice were generated by crossing TAZ flox/flox mice (Jackson Laboratory) with Albumin-Cre mice (Jackson Laboratory). Age and sex-matched knockout and littermate control mice were studied. Mice underwent PTT and RU496 treatment as described in C57BL/6J mice.

### Histology, immunohistochemistry, and plasma insulin and glucagon measurements

Portions of liver were fixed overnight in 4% paraformaldehyde, embedded in paraffin, and sectioned, and the sections generated were hematoxylin and eosin-stained and their histology was assessed. For TAZ immunohistochemistry, liver sections underwent antigen retrieval in boiling 10 mM sodium citrate buffer (pH = 6) for 10 min and then were blocked with 5% goat serum in PBS for 1 h, treated with 3% hydrogen peroxide for 30 min at room temperature, and incubated with primary antibodies overnight at 4°C, followed by horseradish peroxidase (HRP)-conjugated goat anti-rabbit IgG secondary antibodies (Vector Labs) for 1 h at room temperature. Immunoreactivity was visualized by incubating slices with VIP peroxidase substrate (Vector Labs), with cell nuclei being stained blue using methyl green (Sigma). All images were obtained using an EVOS2 microscope (Life Technologies). Plasma insulin and glucagon were measured using commercial chemiluminescent ELISA kits (Alpco).

### Plasmid and adenoviral vector constructs

Human cDNAs of wild-type TAZ and mutants containing deletions of the WW or CC domains (gifts from Dr. Jeff Wrana: Addgene #24809, #24811, and #24816) were cloned into a pcDNA3-FLAG vector by PCR, restriction enzyme digestion, and ligation. A cDNA expressing the human GR was purchased from ATCC and was cloned into a pcDNA3 vector for overexpression in mammalian cells^51^. TAZS89A, TAZS51A, and GR4A mutants were constructed using site-directed mutagenesis (Agilent). The *GRE*-Luc construct was constructed by annealing two complementary DNAs containing three repeats of a canonical GRE (two inverted hexameric half-site motifs separated by a three-base-pair spacer), followed by ligation into a pGL3 basic vector (Agilent). The human *PCK1* and *G6PC* luciferase constructs (*PCK1*-Luc and *G6PC*-Luc) and the plasmids encoding HNF4α and PGC1α were gifts from Dr. Pere Puigserver ^52^. The GFP-GR construct was a gift from Dr. A. Wong (Addgene #47504), the pcDNA3-FLAG-human YAP1 vector was a gift from Dr. Yosef Shaul (Addgene #18881), the *TEADE*-Luc (8x*GTIIC*-Luc) construct containing eight repeats of the TEAD response element was a gift from Dr. Stefano Piccolo (Addgene #34615) ^53^, and the TEAD1 expression vector was a gift from Dr. Kun-Liang Guan (Addgene #33109). The sequences of all the plasmids were confirmed by sequencing.

shRNAs targeting mouse or human TAZ, LacZ, and human lamin were constructed in a U6 Block-it vector (Life Technologies). Adenoviruses expressing these shRNAs were established as per the manufacturer’s instructions (Life Technologies) and as previously reported ^50^. The sequences of the shRNAs are shown in **Supplemental Table 1.** Adenoviruses overexpressing FLAG-tagged human TAZ or control GFP were constructed using the AdTrack system ^54^. All viruses were amplified in 293A cells (Life Technologies), purified using cesium chloride gradient centrifugation, and titered using an end-point dilution method. The 293A cells were maintained in high glucose (4.5 g/L) Dulbecco’s modified Eagle’s medium (DMEM) supplemented with 100 units/ml penicillin, 100 units/ml streptomycin, and 10% FBS at 37°C and in a 5% CO_2_-containing atmosphere in a humidified incubator, and they were free of mycoplasma contamination. Cell culture reagents were purchased from Life Technologies unless otherwise indicated.

### Primary hepatocyte studies

Primary mouse hepatocytes were isolated from Eight- to twelve-week-old male C57BL/6J mice, as described previously, using a two-step collagenase perfusion method ^50^, and they were maintained at 37°C and in a 5% CO_2_-containing atmosphere in a humidified incubator. Hepatocytes were seeded in collagen-coated 6-well dishes at a density of 5 × 10^5^ cells/well in M199 medium supplemented with 2 mM glutamine, 100 units/ml penicillin, 100 units/ml streptomycin, and 10% FBS. After 4 h, the cells were washed with PBS and incubated in fresh medium, in which various treatments were administered.

For adenovirus-mediated overexpression, adenoviruses were added to cell culture media at multiplicity of infection (MOIs) of 5–15 for 24 h prior to harvest. Alternatively, for adenovirus-mediated knockdown, hepatocytes were incubated with adenoviruses at MOIs of 15–50 for 36–48 h prior to harvest. For gene expression studies, on the day of harvest the medium was replaced with fresh M199 medium supplemented with 2 mM glutamine, 100 units/ml penicillin, and 100 units/ml streptomycin. Cells were treated with 20 nM glucagon (Sigma) for 3 h, 100 nM dexamethasone (Sigma) for 6 h, or vehicle. Alternatively, cells were pretreated with RU486 (10 μM) for 30 min prior to glucagon treatment. For immunoblotting studies, cells were incubated overnight in M199 without FBS and then stimulated with insulin (20 nM) for 5 min or glucagon (20 nM) for 30 min. All cells were harvested at the same time, at the conclusion of each experiment.

To measure glucose production, hepatocytes were fasted overnight in low glucose (1 g/L) DMEM media and then incubated in glucose production medium (2 mM L-carnitine and 2 mM pyruvate, without glucose and phenol red) in the presence of glucagon (20 nM). Small aliquots of medium were removed at various time points, and the concentrations of glucose were measured using an Amplex glucose assay kit (Life Technologies), as per the manufacturer’s instructions.

Peripcentral and periportal hepatocytes were isolated as previously described^55^. Briefly, primary mouse hepatocytes isolated via the two-step collagenase perfusion method were suspended in 1 ml DMEM media and subjected to Percoll (GE Healthcare) gradient centrifugation at 1,000g for 30 minutes. The gradient consists of 1 ml 70% Percoll followed by 3 ml 52%, 4 ml 42%, and 5 ml 30% Percoll. The periportal and pericentral cell layers were removed and washed with PBS prior to immunoblotting.

### Luciferase reporter assay

HepG2 cells (HB-8065; ATCC) were maintained in DMEM supplemented with 100 units/ml penicillin, 100 units/ml streptomycin, and 10% FBS at 37°C and in a 5% CO_2_-containing atmosphere, and they were free of mycoplasma contamination. Transient transfection was performed using Lipofectamine 2000 (Life Technologies) as per the manufacturer’s instructions. Briefly, cells were seeded in 24-well dishes on day 0. On day 1, cells were transfected with 50–100 ng/well of the indicated firefly luciferase reporter, 50–100 ng/well of *Renilla* plasmid as an internal control for transfection efficiency, and 10–300 ng/well of expression vector. Empty vector was added to ensure that the same amount of total DNA was used in each transfection reaction. On day 2, the cells were either harvested or placed in serum-free media, with or without 100 nM Dex, for an additional 16–20 h, and were harvested on day 3 by scraping into passive cell lysis buffer (Promega). Luciferase activity was then measured using a commercial dual luciferase assay kit (Promega), after which firefly luciferase activity was normalized to *Renilla* activity and the values for the control groups were set to one.

### RNA isolation and analysis

RNA was isolated using Trizol reagent (Life Technologies), according to the manufacturer’s instructions. For quantitative real-time RT-PCR, 1 μg RNA was reverse transcribed using a High-Capacity cDNA Reverse Transcription Kit (Life Technologies). The resulting cDNA was amplified and quantified using SYBR Green PCR Master Mix (Bioline) in QuantStudio 6 Flex. Real-time qRT-PCR was performed in duplicate or triplicate, and the values obtained for each sample were normalized to the expression of the reference genes TATA box binding protein (*Tbp*) for *in vivo* experiments and ribosomal protein large p0 (*Rplp0*) for *in vitro* experiments. After normalization, the expression in untreated controls was set to one. The sequences of the primers used in real-time qRT-PCR are listed in **Supplemental Table 2**.

### Immunoblotting

Whole cell lysates were prepared by collecting cells in lysis buffer (50 mM Tris, pH 7.5, 150 mM NaCl, 1 mM EDTA, 1% NP-40, 0.5% sodium deoxycholate, 1.0% SDS, 2 mM NaF, and 2 mM Na_3_VO_4_, supplemented with protease and phosphatase inhibitors [Sigma]), sonicating, and then centrifuging at 13,000 × *g* for 10 min ^50^. Some cells were treated with 1 μM DSP (Life Technologies), a cross-linker, for 30 min on ice prior to harvesting. Liver homogenates were prepared by homogenizing liver tissue in lysis buffer ^50^. Nuclear extracts were prepared using an NE-PER extraction kit (Thermo Scientific), according to the manufacturer’s instructions, or as previously described ^56^. Briefly, livers were homogenized in a Dounce homogenizer in hypotonic buffer (15 mM HEPES, pH 7.9, 1.5 mM MgCl_2_, 10 mM KCl, 0.2% NP-40, 1 mM EDTA, 5% sucrose, and 1 mM dithiothreitol, supplemented with protease and phosphatase inhibitors). The homogenate was layered onto a sucrose cushion buffer (300 mM sucrose, 60 mM KCl, 10 mM Tris-HCl, pH 7.5, 1 mM EDTA, 0.15 mM spermine, 0.5 mM spermidine, and 1 mM dithiothreitol) and centrifuged at 2,000 × *g* for 2 min.

Lysates were subjected to SDS-PAGE and transferred onto a PVDF membrane (Thermo Scientific). After blocking, blots were incubated overnight with primary antibody (1:1,000 to 5,000 dilution), and then secondary antibody conjugated to horseradish peroxidase (Thermo Scientific) and chemiluminescent ECL reagents (Thermo Scientific) were used to identify specific bands. Band intensities were determined using ImageJ and normalized to the intensity of loading control bands and the values of controls were set to 1. The antibodies used in this study were obtained from commercial sources and are listed in **Supplemental Table 3**.

### Co- immunoprecipitation

Because of poor transfection efficiency in HepG2 and primary mouse hepatocytes, co-IP assays were performed in 293A cells after the co-transfection of expression vectors, as indicated in the Figures, for 36–48 h. Cells were lysed in RIPA buffer (50 mM Tris, pH 7.5, 150 mM NaCl, 1 mM EDTA, 0.5% NP40, and 0.5% sodium deoxycholate) supplemented with protease and phosphatase inhibitors, sonicated, and centrifuged at 13,000 × *g* for 10 min. The resulting cell lysates were incubated with anti-FLAG antibody overnight at 4°C. Immunoprecipitates were obtained by adding Protein A/G agarose (Santa Cruz Biotechnologies), washed three times with RIPA buffer containing 0.025–0.05% SDS, and subjected to immunoblotting. Alternatively, IP was performed using nuclear extracts from mouse livers. The antibodies used for IP are listed in **Supplemental Table 3**.

### Chromatin immunoprecipitation

ChIP assays using primary hepatocytes and liver tissues were performed as previously described, with minor modifications ^23, 56^. Cells were cross-linked with 1% formaldehyde at room temperature for 15 min, and then the reaction was stopped by the addition of 125 mM glycine for 5 min at room temperature. Cells were washed with PBS, collected in harvest buffer (100 mM Tris-HCl, pH 9.4, and 10 mM DTT), incubated on ice for 10 min, and then centrifuged at 2,000 × *g* for 5 min. The cells were then sequentially washed with ice-cold PBS, buffer I (0.25% Triton X-100, 10 mM EDTA, 0.5 mM EGTA, and 10 mM HEPES, pH 6.5), and buffer II (200 mM NaCl, 1 mM EDTA, 0.5 mM EGTA, and 10 mM HEPES, pH 6.5). Livers were prepared in a similar manner: livers from two to four mice were pooled, minced into small pieces, cross-linked with 1% formaldehyde, and homogenized in hypotonic buffer (15 mM HEPES, pH 7.9, 1.5 mM MgCl_2_, 10 mM KCl, 0.2% NP-40, 1 mM EDTA, 5% sucrose, and 1 mM dithiothreitol) supplemented with protease and phosphatase inhibitor cocktails. The nuclei were isolated by centrifugation of the resulting homogenates laid over a cushion buffer (300 mM sucrose, 60 mM KCl, 10 mM Tris-HCl, pH 7.5, and 1 mM EDTA). The pellets were resuspended in lysis buffer (1% SDS, 10 mM EDTA, and 50 mM Tris-HCl, pH 8.0) supplemented with protease and phosphatase inhibitor cocktails; sonicated to reduce the DNA length to 0.3–1.5 kb; and then centrifuged at 8,000 × *g* for 1 min at 4^°^C. The soluble chromatin was diluted 10-fold in dilution buffer (1% Triton X-100, 2 mM EDTA, 150 mM NaCl, and 20 mM Tris-HCl, pH 8.0) and then pre-cleared with protein A/G sepharose beads (Santa Cruz Biotechnology) at 4°C for 1 h (protein A/G sepharose was incubated with sheared salmon sperm DNA and washed three times in dilution buffer prior to use). Equal amounts of pre-cleared chromatin were then added to 2–3 μg of antibody overnight. The next day, 25 μl protein A/G sepharose beads were added and the incubation was continued for another 2 h. The beads were collected and then washed in TSE I (0.1% SDS, 1% Triton X-100, 2 mM EDTA, 20 mM Tris-HCl, pH 8.0, and 150 mM NaCl), TSE II (0.1% SDS, 1% Triton X-100, 2 mM EDTA, 20 mM Tris-HCl, pH 8.1, and 500 mM NaCl), and TE buffer (10 mM Tris, pH 8.0, and 1 mM EDTA), and were finally eluted in 1% SDS/0.1 M NaHCO_3_. Eluates were heated to 65°C for at least 6 h to reverse the formaldehyde cross-linking and then treated with proteinase K (Life Technologies). DNA fragments were isolated using a DNA purification kit (Qiagen). The immunoprecipitated DNA and 10% of the pre-cleared chromatin (input DNA) were then subjected to real-time PCR using Power SYBR Green PCR Master Mix (Biolines), in triplicate. For each immunoprecipitate, we calculated the relative enrichment as 2^−ΔCt^, where ΔCt was calculated as the mean Ct value obtained from the immunoprecipitate DNA minus the mean Ct value obtained from the input DNA. The mean relative enrichment of the replicate immunoprecipitates (3–4 per group) was calculated, and the results are expressed in arbitrary units, with the IgG control for each primer pair set to one. Representative results from two to four independent experiments are shown. The antibodies used in the ChIP assays are listed in **Supplemental Table 3** and the real-time PCR primers are listed in **Supplemental Table 2**.

## Statistics

Mice were randomized into treatment groups and sample numbers are indicated in figure legends. Data are presented as mean and SEMs of biological repeats. Representative results of two to five independent experiments are shown. Data were analyzed by two-tailed unpaired Student’s *t*-test or ANOVA for repeated measurements with multiple comparison post-hoc test using Graphpad Prism software.

## Supplemental Information

### Supplemental Figures and Legends

**Figure 1–Figure supplement 1.**
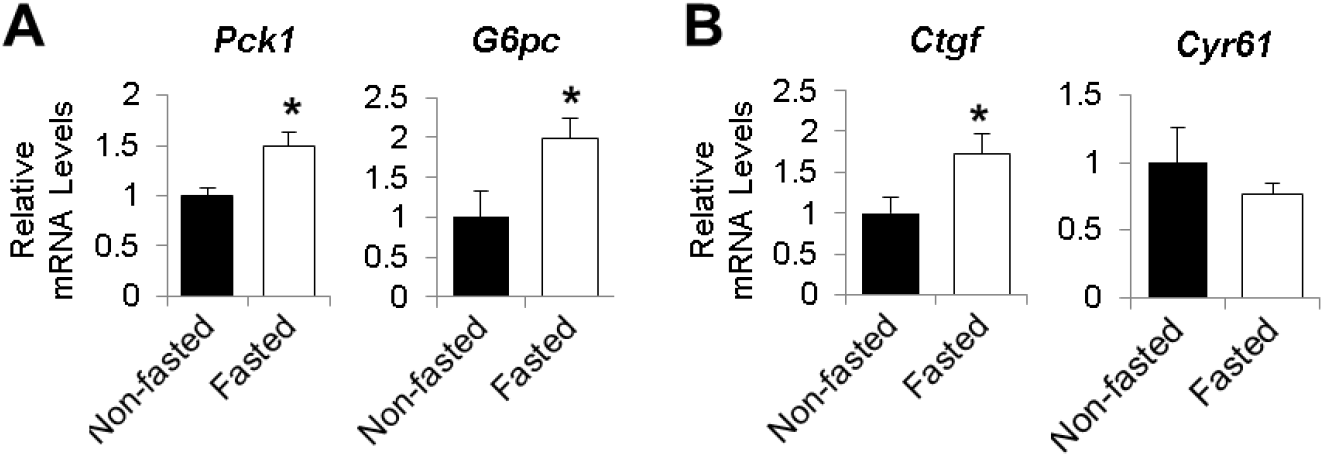
**Figure supplement 1.** Hepatic gene expression in fed and fasted mouse livers. Eight- to twelve-week-old C57BL/6J mice were fed *ad libitum* (Non-fasted) or fasted for 24 h (Fasted). Gene expression was measured by real-time qRT-PCR. Data are means and SEMs; controls values were set to 1; n = 6; \**p* < 0.05 (unpaired Student’s *t*-test).

**Figure 1–Figure supplement 2.**
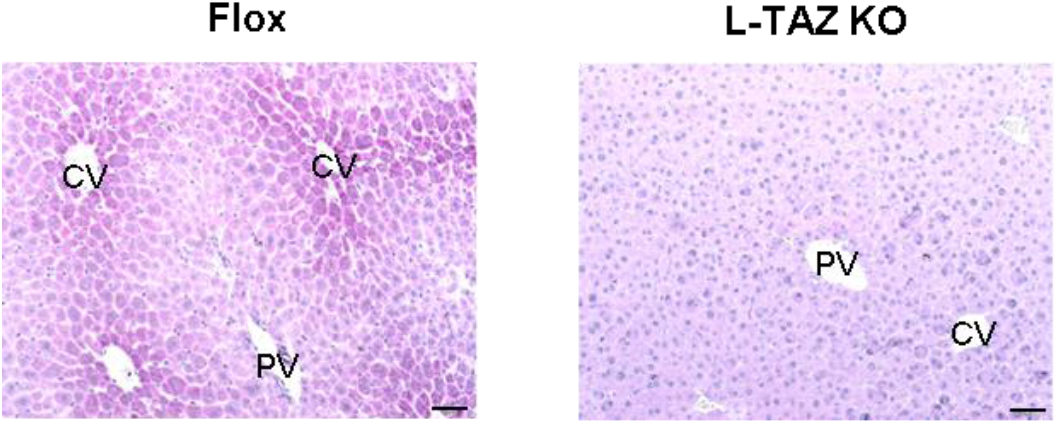
**Figure supplement 2.** Validation of immunohistochemical staining for TAZ in control and L-TAZ KO liver. Immunohistochemistry using an anti-TAZ antibody in L-TAZ KO and control floxed (Flox) mouse liver sections (CV, central vein; PV, portal vein; scale bar, 50 μM).

**Figure 1 – Figure supplement 3.**
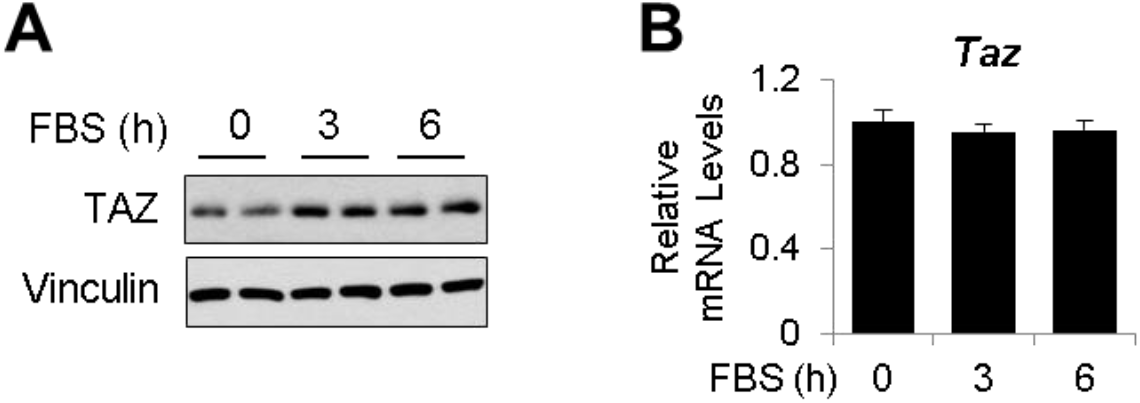
**Figure supplement 3.** mRNA and protein expression of TAZ in primary mouse hepatocytes. Primary mouse hepatocytes were isolated from eight- to twelve-week-old C57BL/6J male mice. Cells were incubated in low glucose media (1g/L) without FBS for overnight and then placed in high glucose media (4.5g/L) with FBS for the indicated times. Protein levels (A) and mRNA expression (B) were measured. Data are means and SEMs of triplicated wells. Data were analyzed by one-way ANOVA.

**Figure 2–Figure supplement 1.**
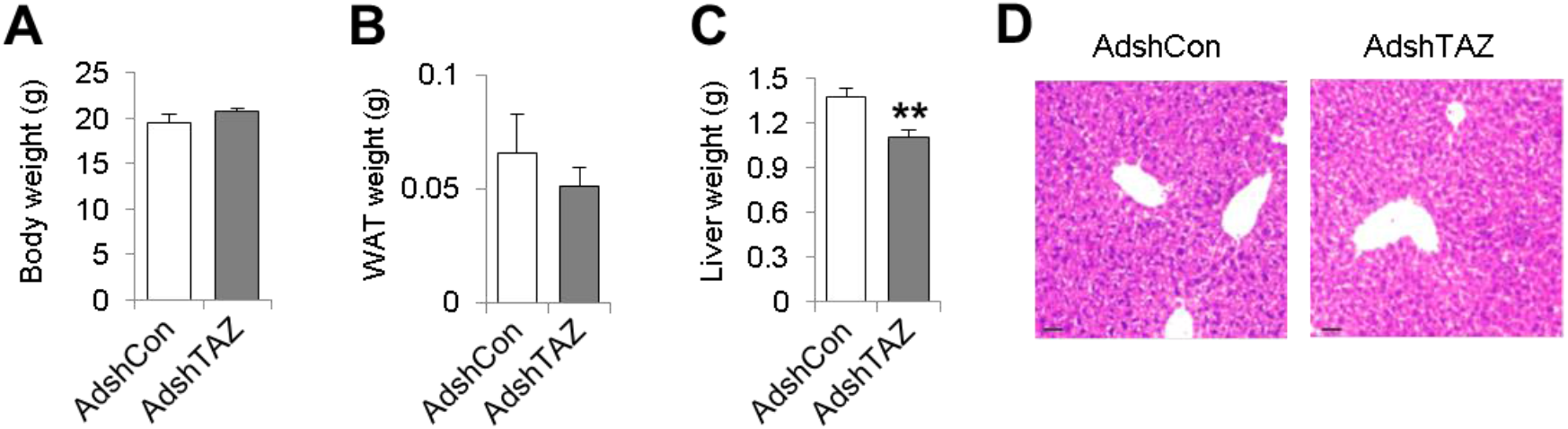
**Figure supplement 1.** Information on mice administered AdshTAZ or AdshCon. Eight- to twelve-week-old C57BL/6J mice were administered AdshTAZ or AdshCon, then sacrificed 8 days later in the *ad libitum*-fed state. Body (A), epididymal white adipose tissue (WAT) (B), and liver weight (C) were measured. (D) H and E staining of liver sections (scale bar, 50μM). Data are means and SEMs; control values were set to 1; n = 7–9. Data were analyzed by unpaired Student’s *t*-test; ** *p*<0.01.

**Figure 2–Figure supplement 2.**
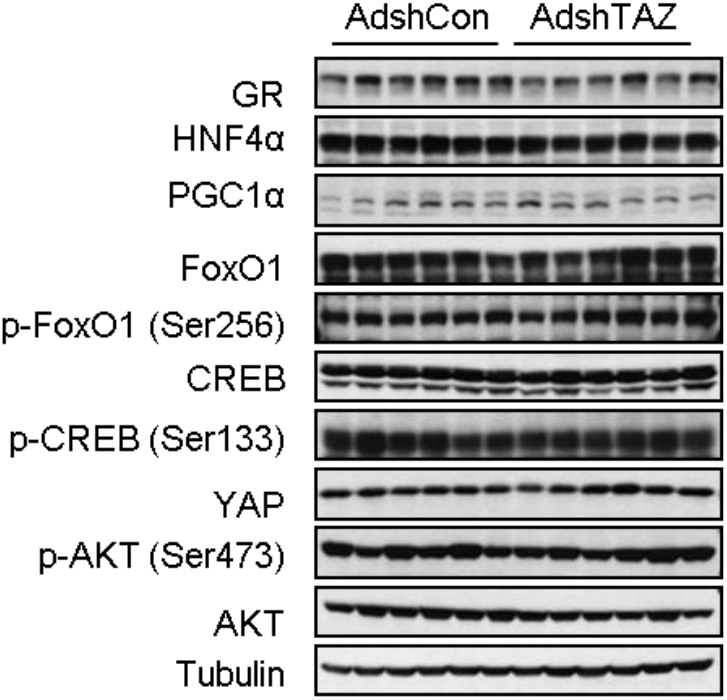
**Figure supplement 2.** TAZ knockdown has no effects on protein expression of key gluconeogenic factors and insulin and glucagon signaling in mice. Eight- to twelve-week-old C57BL/6J mice were administered AdshTAZ or AdshCon, then sacrificed 8 days later in the *ad libitum*-fed state. Hepatic proteins were measured by immunoblotting whole cell lysates.

**Figure 2–Figure supplement 3.**
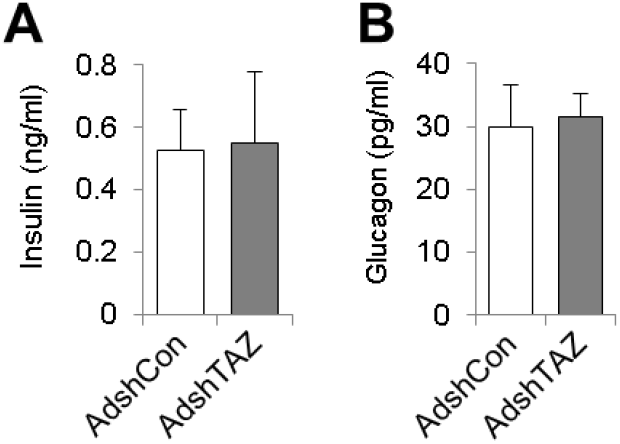
**Figure supplement 3.** Hepatic TAZ knockdown has no effects on plasma insulin and glucagon concentrations. Eight- to twelve-week-old C57BL/6J mice were administered AdshTAZ or AdshCon, then sacrificed 8 days later in the *ad libitum*-fed state. Plasma insulin (F) and glucagon (G) were measured. Data are means and SEMs; n = 7–9. Data were analyzed by unpaired Student’s *t*-test.

**Figure 2–Figure supplement 4.**
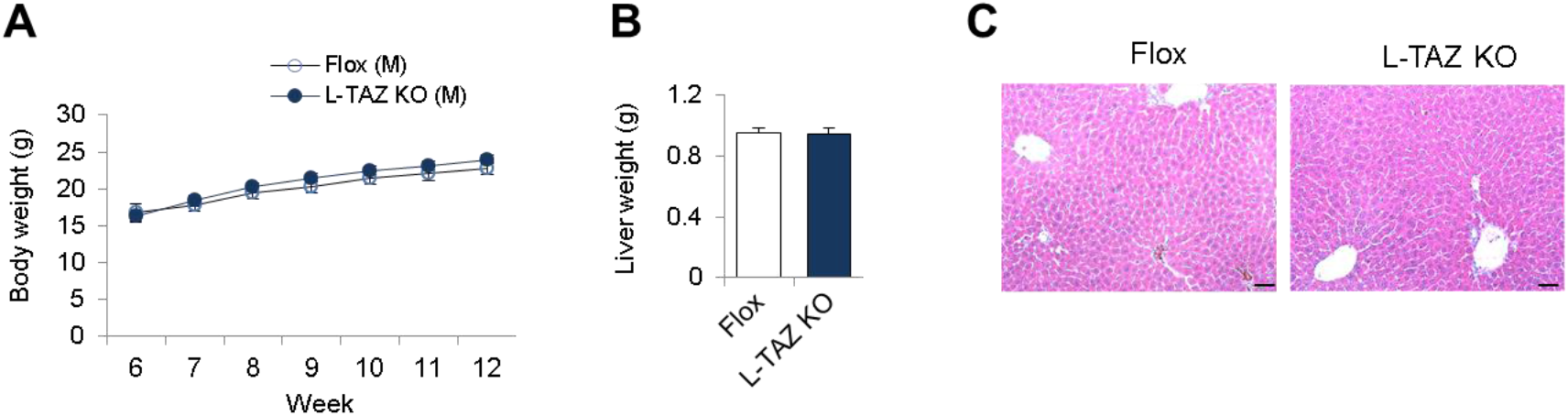
**Figure supplement 4.** Information on L-TAZ KO and flox mice. Age and sex-matched male L-TAZ KO and littermate controls were studied. Body (A) and liver weight (B) were measured. (C) H and E staining of liver sections (scale bar, 50μM). Data are means and SEMs; control values were set to 1; n = 7–8. Data were analyzed by two-way ANOVA (A) and unpaired Student’s *t*-test (B).

**Figure 2–Figure supplement 5.**
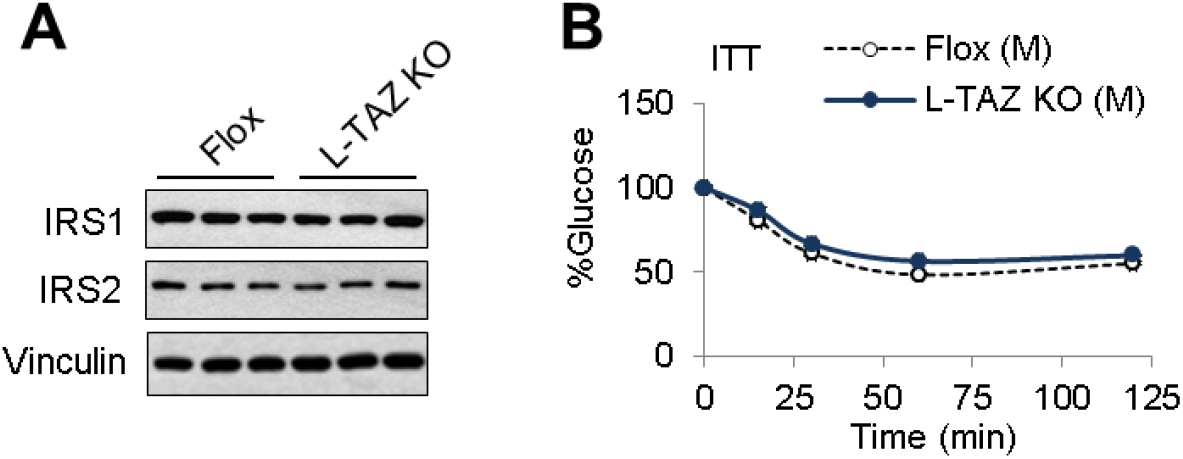
**Figure supplement 5.** L-TAZ KO has no effects on insulin sensitivity. Eight- to twelve-week-old age and sex-matched male L-TAZ KO and littermate controls were studied. (A) Hepatic proteins were measured by immunoblotting whole cell lysates. (B) Insulin sensitivity was assessed by intraperitoneal insulin tolerance test (ITT). Data are means and SEMs; n = 7–8. Data were analyzed by two-way ANOVA.

**Figure 3–Figure supplement 1.**
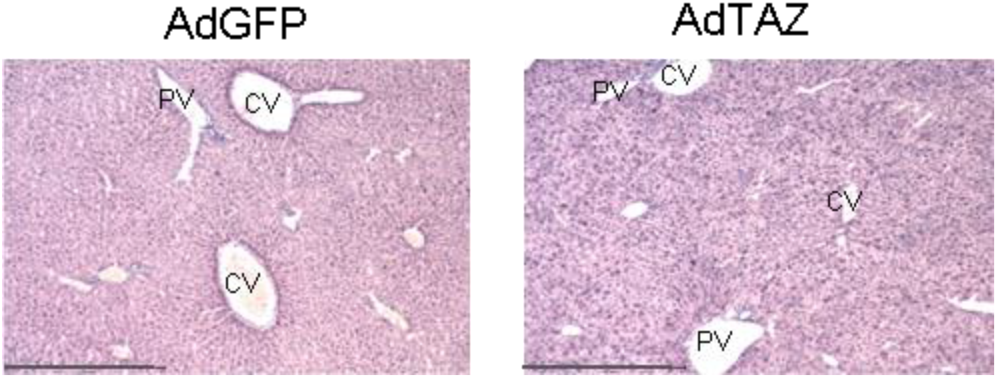
**Figure supplement 1.** Immunohistochemical staining for TAZ in livers of mice administered AdTAZ or control AdGFP. Eight- to twelve-week-old C57BL/6J mice were administered AdGFP or AdTAZ, then sacrificed 5 days later, after a 24-h fast. CV, central vein; PV, portal vein; scale bar, 500 μM.

**Figure 3–Figure supplement 2.**
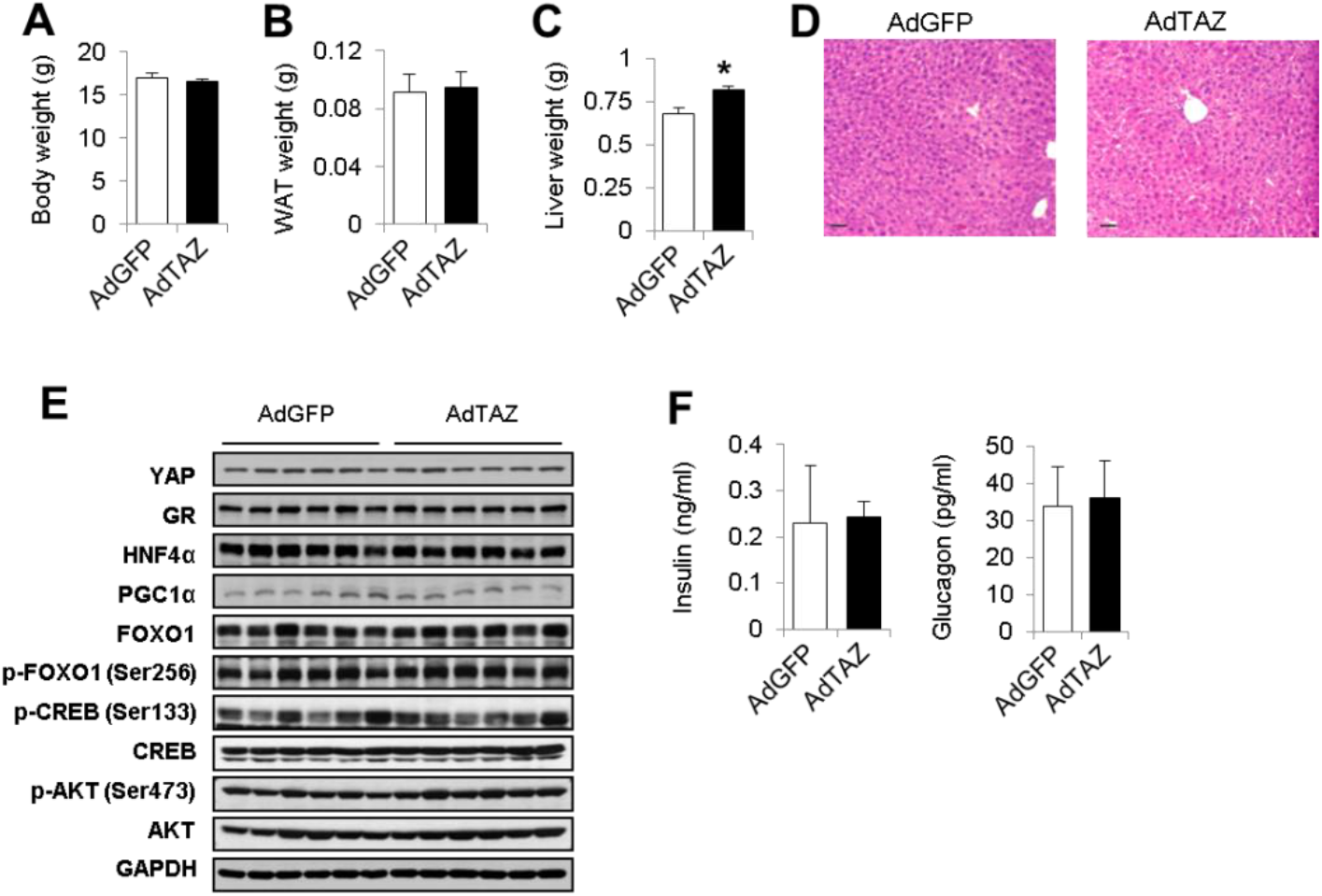
**Figure supplement 2.** Information on mice administered AdTAZ or control AdGFP Eight- to twelve-week-old C57BL/6J mice were administered AdGFP or AdTAZ, then sacrificed 5 days later, after a 24-h fast. Mouse body weight (A), epididymal white adipose tissue weight (WAT) weight, (B), liver weight (C), and plasma insulin (F, left) and glucagon (F, right) levels were measured. (D) H and E staining of liver sections (scale bar, 50μM). (E) Hepatic proteins were measured by immunoblotting whole cell lysates. Data are means and SEMs; controls values were set to 1; n = 5-10. Data were analyzed by unpaired Student’s *t*-test; \**p* < 0.05.

**Figure 4–Figure supplement 1.**
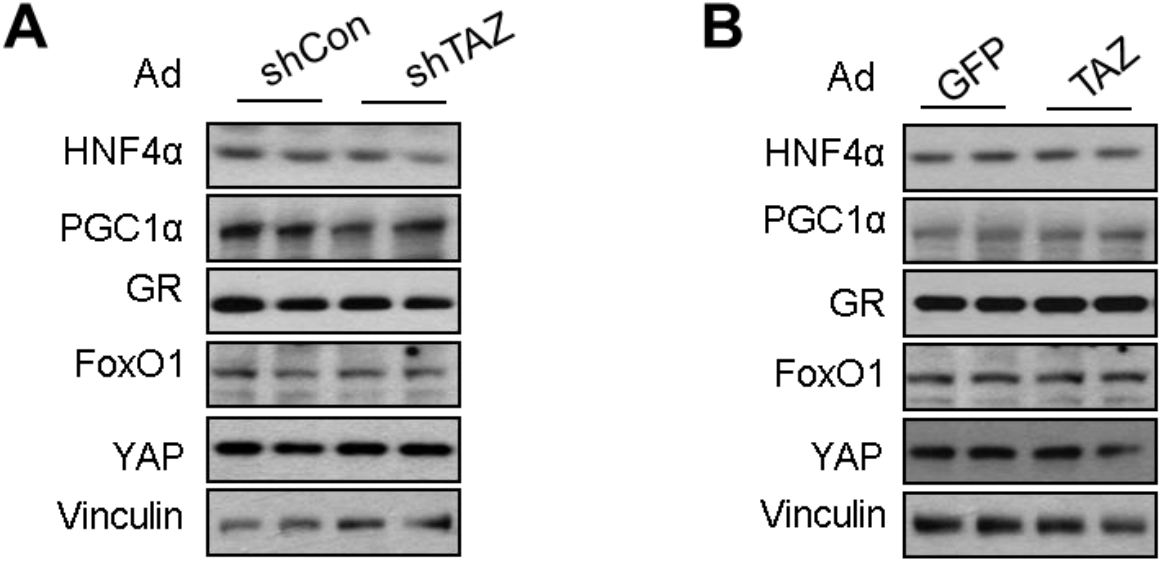
**Figure supplement 1.** Protein expression in primary mouse hepatocytes in the presence of TAZ knockdown or overexpression. Primary mouse hepatocytes were isolated from Eight- to twelve-week-old C57BL/6J male mice. (A) To knockdown TAZ, cells were infected with AdshTAZ or AdshCon. (B) To overexpress TAZ, cells were infected with AdTAZ or AdGFP. Protein levels were measured by immunoblotting whole cell lysates.

**Figure 4–Figure supplement 2.**
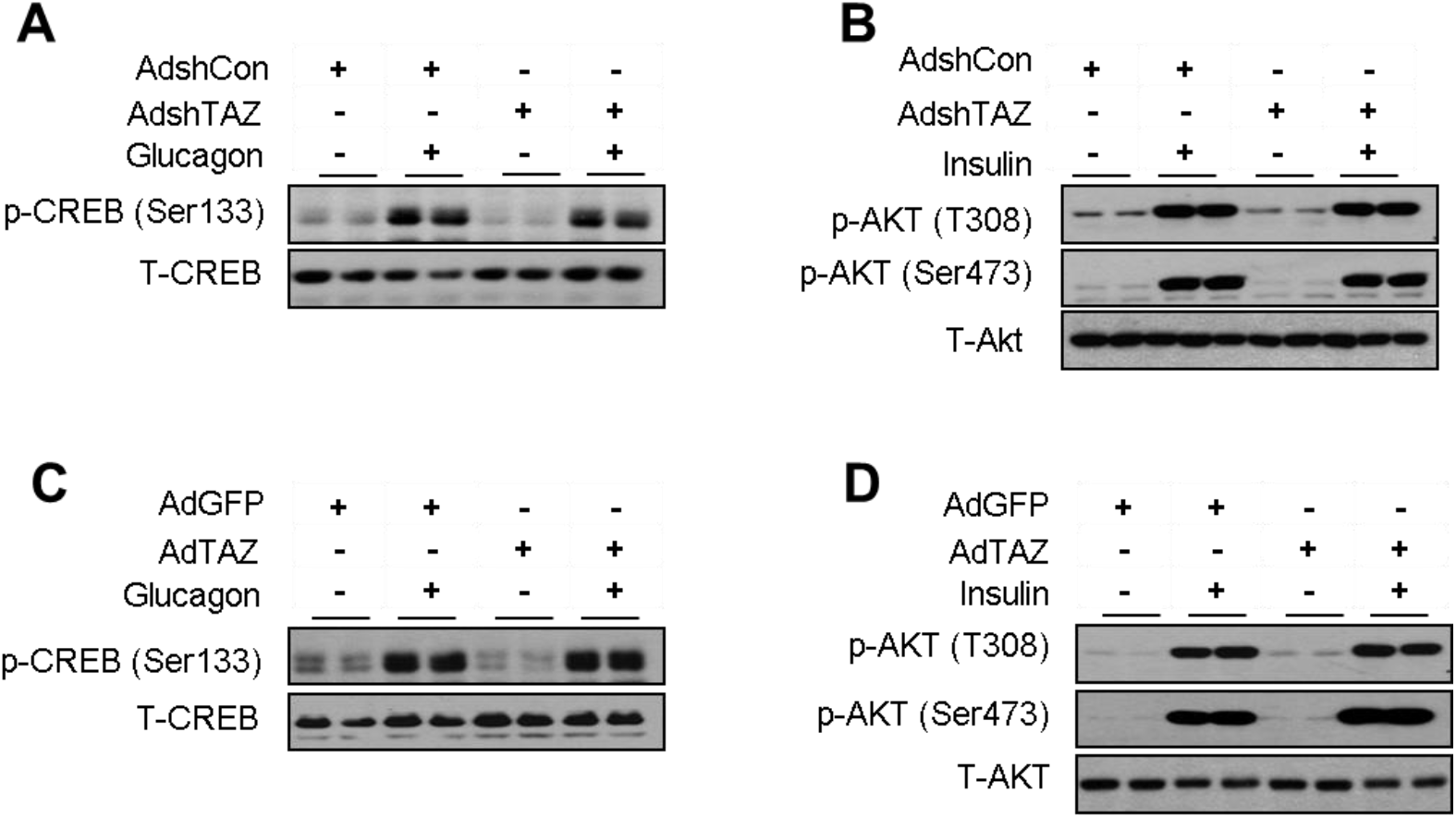
**Figure supplement 2.** TAZ knockdown or overexpression in primary hepatocytes has no effects on insulin or glucagon signaling. Primary mouse hepatocytes were infected with AdshTAZ or AdshCon (A–B) or AdTAZ or AdGFP (C–D). Protein levels were measured by immunoblotting whole cell lysates. Cells were treated with glucagon (20 nM) for 30 min (A and C) or insulin (20 nM) for 10 min (B and D). Protein levels were measured by immunoblotting whole cell lysates.

**Figure 5–Figure supplement 1.**
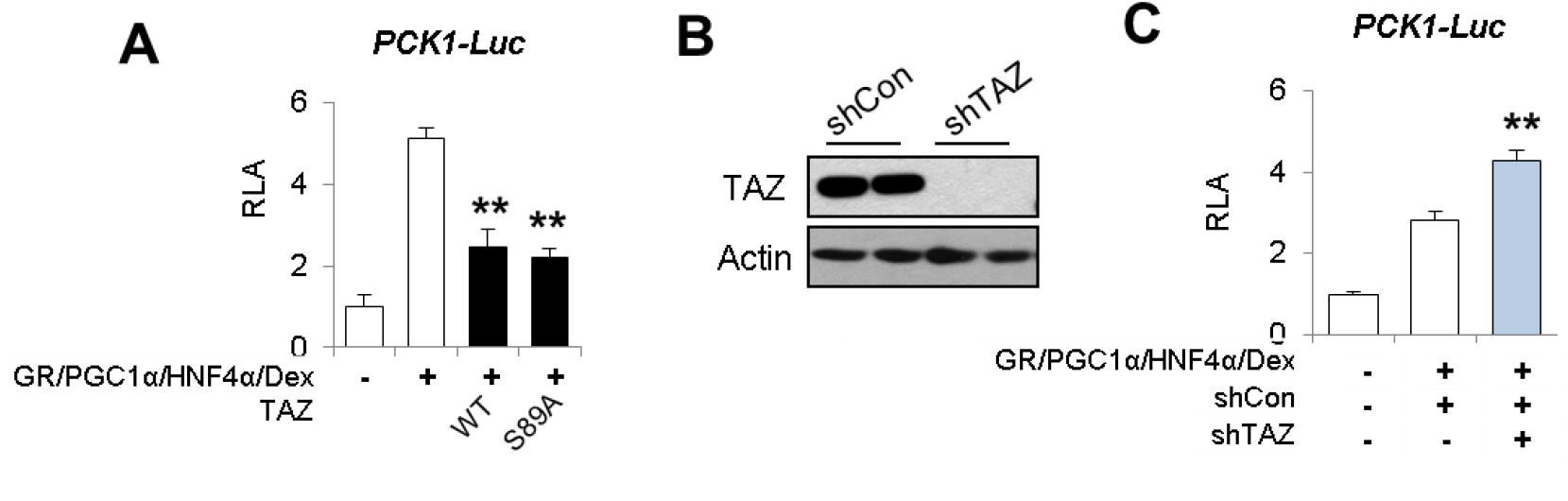
**Figure supplement 1.** TAZ inhibits PCK-Luc activity. (A and C) HepG2 cells were co-transfected with expression vectors, luciferase reporters, and an internal control (*Renilla*), and treated with dexamethasone (Dex) (100 nM), as indicated. Relative Luciferase Activity (RLA) is presented after normalization to the *Renilla* activity. (B) Cells were co-transfected with a Flag-TAZ expression vector and a TAZ knockdown vector, or a control vector, as indicated and TAZ expression was measured by immunoblotting whole cell lysates. Data are means and SEMs of triplicated wells; controls were set to 1.Data were analyzed by one-way ANOVA. \*\**p* < 0.01 denotes comparisons with wells treated with GR/PGC1α/HNF4α/Dex alone.

**Figure 5–Figure supplement 2.**
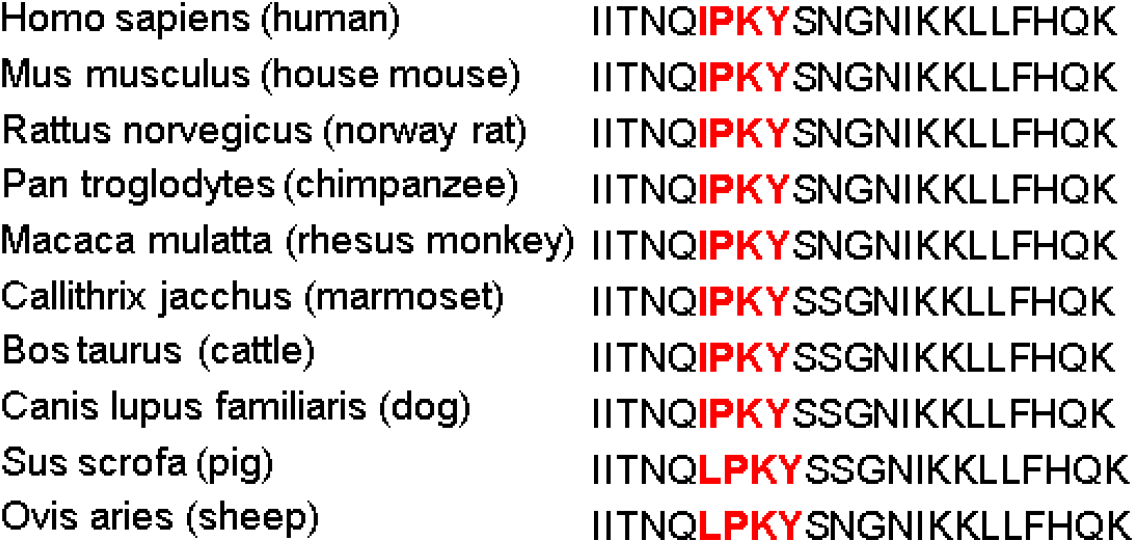
**Figure supplement 2.** A conserved I/LPKY motif is identified in GR from various species. Highlighted in **red**.

**Figure 5–Figure supplement 3.**
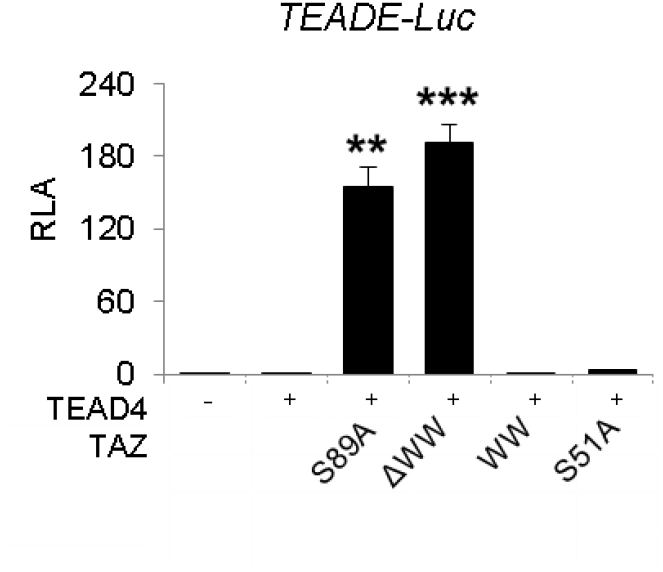
**Figure supplement 3.** Effects of TAZ mutants on TEAD-transactivation. 293A cells were co-transfected with expression vectors, luciferase reporters, and an internal control (*Renilla*), and treated with dexamethasone (Dex) (100 nM), as indicated. Relative Luciferase Activity (RLA) is presented after normalization to the *Renilla* activity. Data are means and SEMs of triplicated wells; controls were set to 1. Data were analyzed by one-way ANOVA. \*\**p* < 0.01 and \*\*\**p* < 0.001 denotes comparisons with wells transfected with TEAD4 alone.

**Figure 5–Figure supplement 4.**
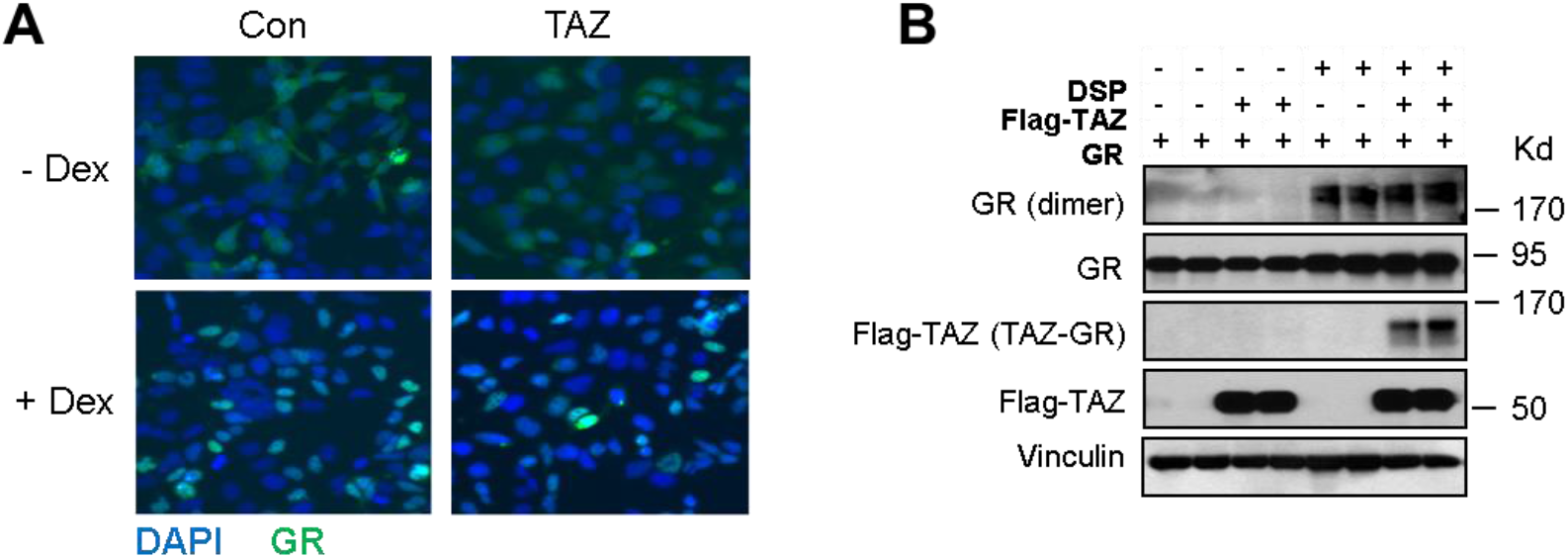
**Figure supplement 4.** TAZ has no effects on GR nuclear localization or the amounts of GR dimer in cells. (A) 293A cells were transfected with expression vectors for GR fused with a GFP, TAZ, or a control empty vector (Con), and treated with Dex (100 nM) or vehicle for 6 h prior to fixation and DAPI staining. GFP-GR distribution was examined by a fluorescent microscope. (B) 293A cells were transfected with expression vectors for GR, TAZ or a control empty vector for 48 hours. Cells were then treated with DSP (3 μM) for 30 minutes on ice and GR dimer was measured by immunoblotting whole cell lysates.

**Figure 6–Figure supplement 1.**
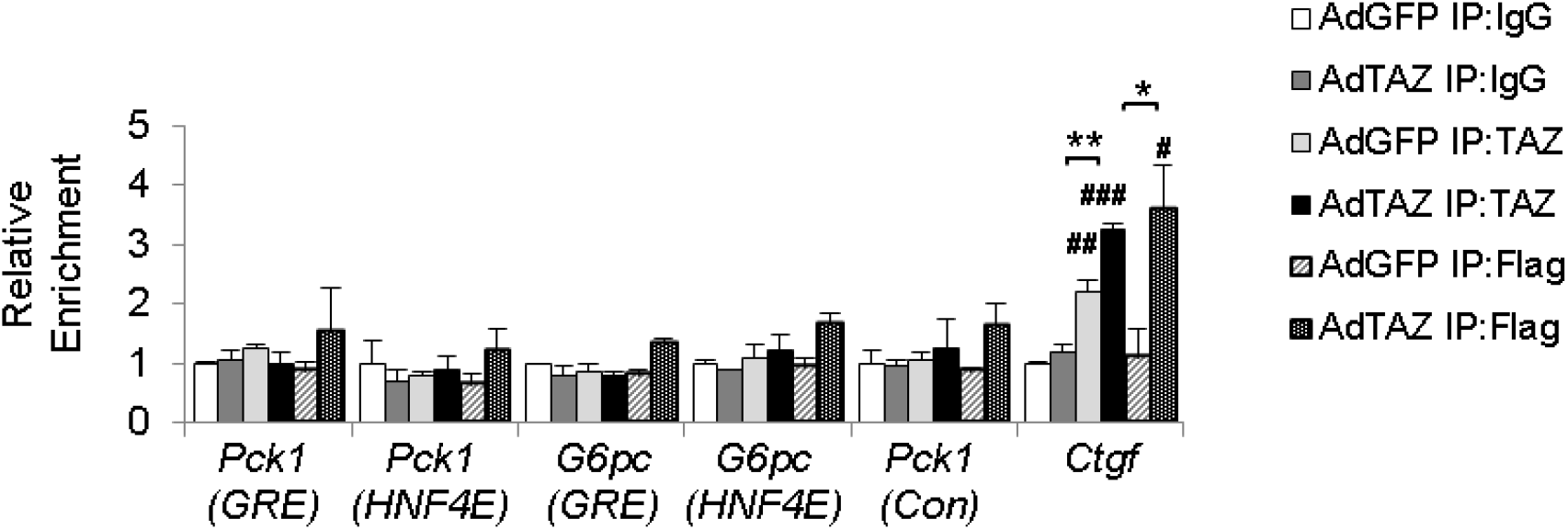
**Figure supplement 1.** TAZ does not bind to gluconeogenic gene promoters. Eight- to twelve-week-old male C57BL/6J mice were administered adenoviruses expressing TAZ (flag-tagged) or GFP for 5 days. ChIP assays were performed from liver extracts using indicated antibodies. Data are means and SEMs of 3–4 immunoprecipitates. Data were analyzed by two-way ANOVA; * *p* < 0.05 and ** *p* < 0.01; # *p* < 0.05, ## *p* < 0.01, and ### *p* < 0.001; # denotes a comparison with IgG.

**Figure 7–Figure supplement 1.**
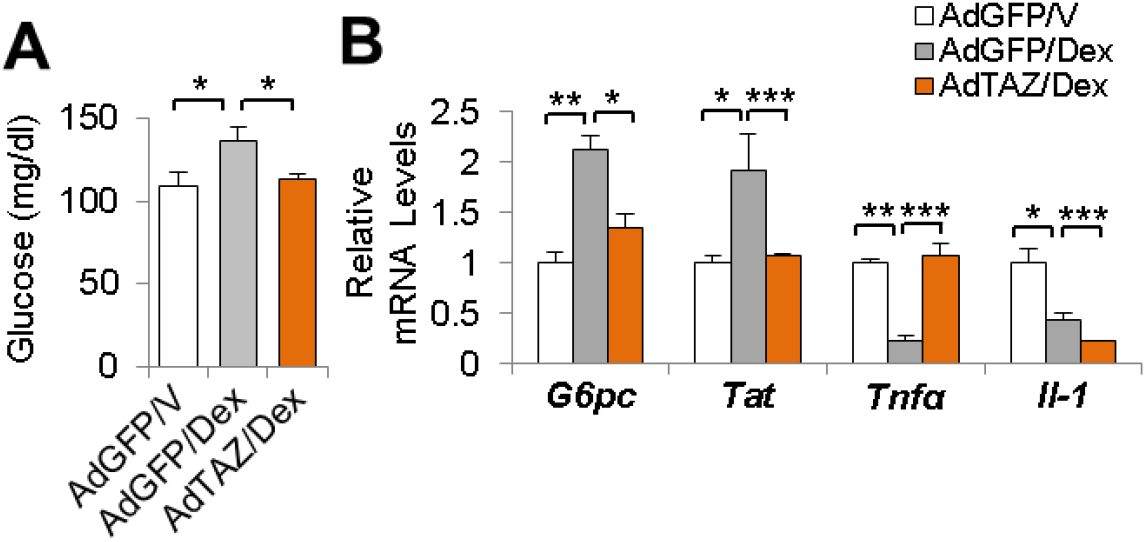
**Figure supplement 1.** Blood glucose concentrations and hepatic gene expression of mice administered AdTAZ and Dex. Eight- to twelve-week-old C57BL/6J mice were administered adenoviruses expressing TAZ or GFP for 5 days. Mice were treated with Dex or vehicle. Blood glucose (A) and hepatic gene expression (B) were measured. Data are means and SEMs; n=5–7. Data were analyzed by one-way ANOVA; * *p* < 0.05, \*\**p* < 0.01, and *** *p* < 0.001.

**Figure 7–Figure supplement 2.**
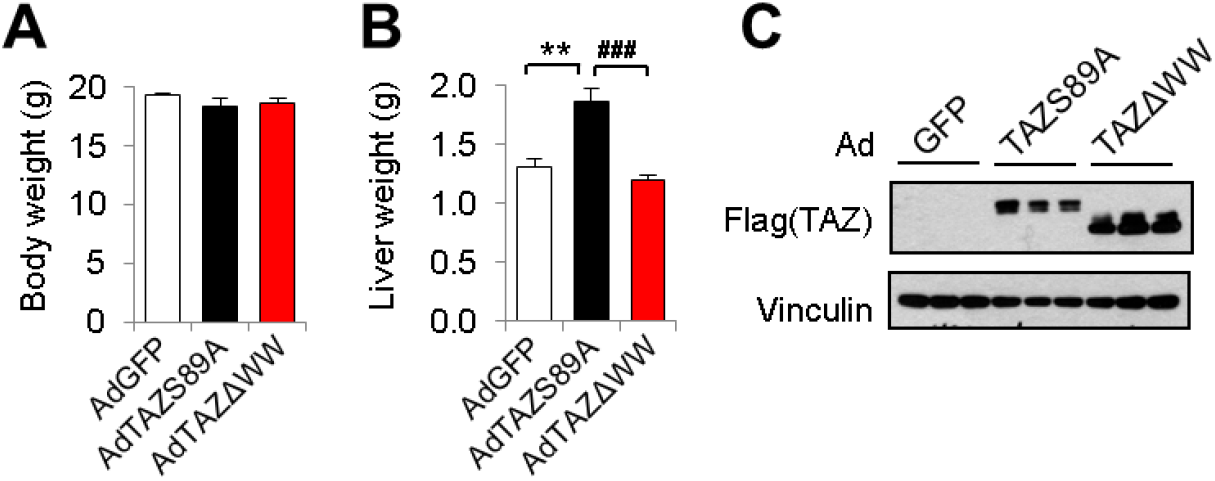
**Figure supplement 2.** Body and liver mass and hepatic TAZ expression in mice administered AdTAZ mutants or AdGFP. Eight- to twelve-week-old C57BL/6J mice were administered adenoviruses expressing TAZ mutants (flag-tagged) or GFP for 5 days. Body weight (A) and liver weight (B) were measured and hepatic protein levels were measured by immunoblotting whole cell lysates. Data are means and SEMs; n = 7–8. Data were analyzed by one-way ANOVA; \*\**p* < 0.01 and ### *p* < 0.001.

**Figure 7–Figure supplement 3.**
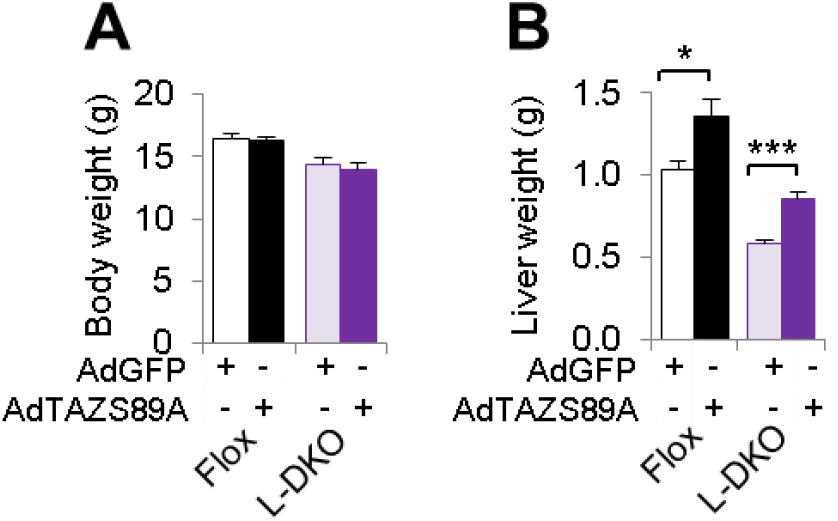
**Figure supplement 3.** Body and liver mass of L-DKO and flox mice treated with AdTAZS89A. Eight- to twelve-week-old female L-DKO and flox controls were administered adenoviruses expressing TAZ or GFP for 5 days. Body (A) and liver (B) weigh were measured. Data are means and SEMs; n = 7–8. Data were analyzed by unpaired Student’s *t*-test \**p* < 0.05 and \*\*\**p* < 0.001.

### Supplemental Tables

**Supplemental Table 1.**
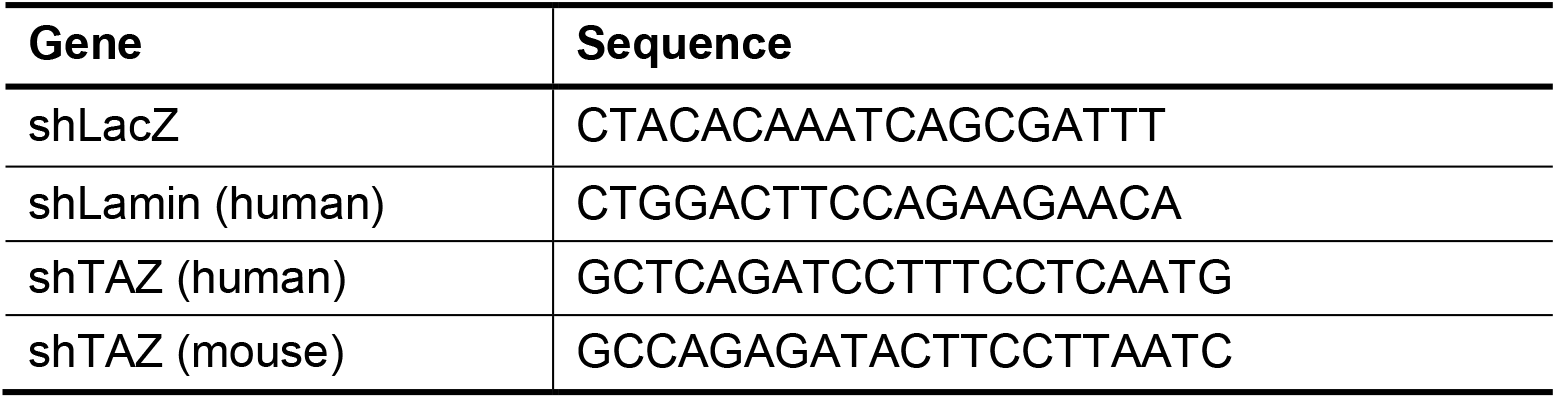
shRNA sequences.

**Supplemental Table 2.**
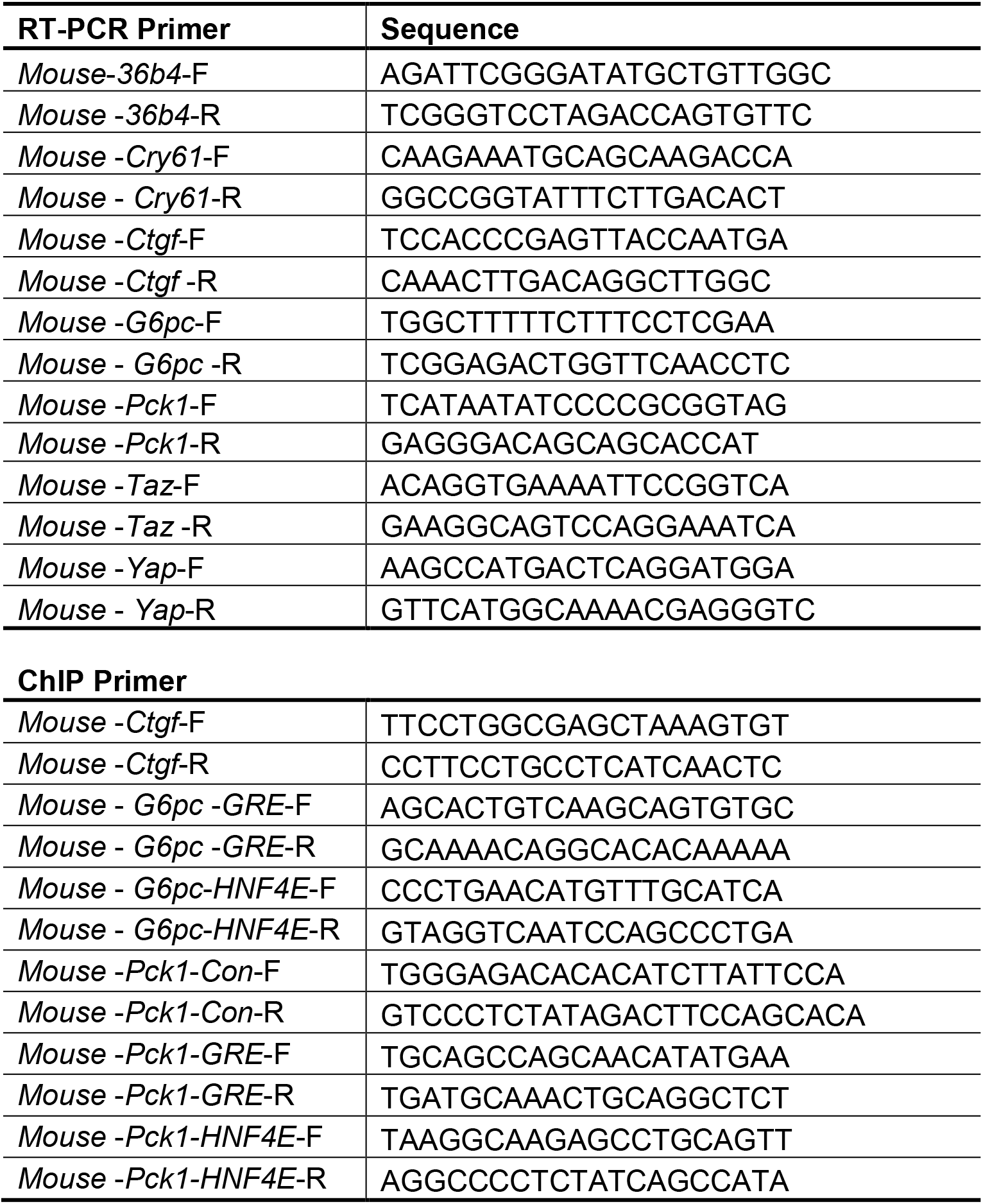
Real-time RT-PCR and ChIP primer sequences.

**Table 3.**
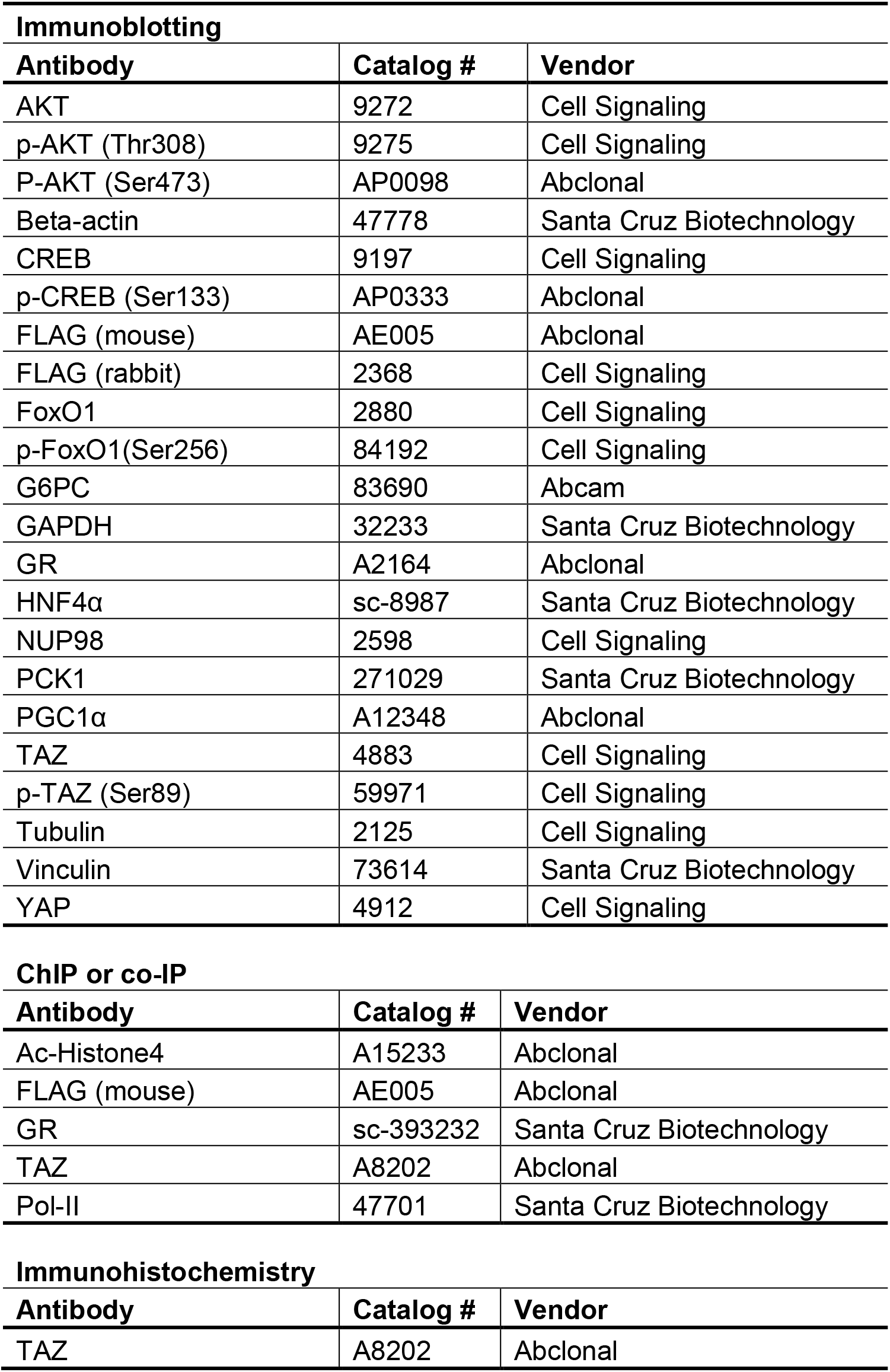
Supplemental Antibodies used in this study.

